# Angiogenic CD8 T cells from PWH induce Granzymes-dependent PAR1 activation promoting endothelial inflammation

**DOI:** 10.1101/2025.08.06.668764

**Authors:** Tong Li, Chinmayee Mehta, Cooper James, Jinya Chen, Hui Chen, Gerard P Ahern, Ziang Zhu, Maitane Faus Cid, Maryam Abdussamad, Shiwen Yu, Jai Kumar, Princy Kumar, Colleen Hadigan, Kyle DiVito, Dorian B McGavern, Leonid Pobezinsky, Marta Catalfamo

**Affiliations:** Department of Microbiology and Immunology, Georgetown University School of Medicine. Washington DC, USA; Department of Pharmacology and Physiology, Georgetown University School of Medicine. Washington DC, USA; Division of Infectious Diseases and Travel Medicine, Georgetown University School of Medicine. Washington DC, USA; Laboratory of Immunoregulation, National Institute of Allergy and Infectious Diseases, National Institutes of Health, Bethesda, MD 20892, USA; Department of Biochemistry, Molecular and Cellular Biology, Georgetown University School of Medicine. Washington DC, USA; Viral Immunology and Intravital Imaging Section, National Institute of Neurological Disorders and Stroke, National Institutes of Health, Bethesda, Maryland, USA; Department of Veterinary and Animal Science, University of Massachusetts, Amherst, MA, USA

## Abstract

In people with HIV (PWH), T cell immune activation and endothelial inflammation are contributors of the increased cardiovascular risk, however the mechanisms remain poorly understood. PAR1 connects the coagulation cascade, endothelial cells and CD8 T cells at the site of endothelial inflammation and we hypothesized that HIV driven CD8 T cell immune activation alters endothelial repair mechanisms. Endothelial repair is partially mediated by angiogenic T (T_ang_) cells that facilitate proliferation of endothelial cells, and differentiation of endothelial progenitor cells at site of vascular injury. During LCMV infection, we identified a subset of CD31^high^CD8 T cells that exhibited a long-lived memory precursor phenotype (CD127^+^KLRG1^-^) and secreted the proangiogenic cytokine vascular endothelial growth factor (VEGF) after viral control. Furthermore, in PWH, the frequencies of CD8 T_ang_ cells were reduced and showed an activated phenotype and expression of granzymes. GZMA^+^GZMB^+^CD8 T_ang_ cells correlated with atherosclerotic cardiovascular disease (ASCVD) risk. *In vitro*, granzyme dependent PAR1 activation led to calcium mobilization and secretion of proinflammatory cytokines IL-6, IL-8 and angiopoietin-2 by primary human endothelial cells. Altogether, these findings suggest that CD8 T cells are involved in immunity against viruses and endothelial homeostasis and HIV driven immune activation alters these functions.

## Introduction

Immune activation remains the hallmark of HIV infection, despite successful suppression of viremia by antiretroviral therapy (ART). HIV-associated inflammation and immune activation drives cardiovascular risk, independent of traditional risk factors, duration of antiretroviral treatment and CD4 counts (1–8). Compared to the general population, People with HIV (PWH) have twice the risk of cardiovascular disease (9–11). The molecular pathways by which HIV drives cardiovascular risk are largely unknown and multifactorial including chronic immune activation, HIV persistence and translation of viral proteins, coinfections, and others (12–21). In PWH, CD8 T cell immune activation is associated with carotid intima-media thickness and coronary plaque suggesting an interplay between CD8 T cell activation and endothelial inflammation and dysfunction (6, 16–18, 22).

During systemic infections such as HIV, endothelial activation and injury can be mediated by viral proteins as well as the host response (23, 24). We hypothesized that HIV driven CD8 T cell immune activation promotes endothelial inflammation altering repair mechanisms. Endothelial repair, is mediated in part by angiogenic T cells (T_ang_) that support proliferation and differentiation of endothelial progenitor cells/endothelial colony forming cells (EPC/ECFC) at the site of endothelial injury (25–27). T_ang_ cells were originally described in the center of endothelial progenitor cell colonies isolated from human peripheral mononuclear cells (PBMCs)(25). In *in vivo* experiments in mice CD31^+^ T_ang_ cells support new vessel formation (25). In inflammatory diseases and aging, reduced frequencies of T_ang_ cells and loss of function have been associated with increased cardiovascular disease risk (25, 28, 29). However, their role in the setting of viral infections remains largely unknown.

The Protease Activated Receptor 1 (PAR1) mediates the crosstalk between the coagulation system, endothelium and the adaptive immune system (CD8 T cells) at sites of vascular inflammation (30–33). PAR1 is one of the four members of the seven transmembrane domain G protein-coupled receptor family (PAR1, PAR2, PAR3 and PAR4) (34). PAR’s activation is mediated by proteolytic cleavage of the N-terminus that uncovers a tethered ligand (35, 36). In humans, PAR1, PAR3 and PAR4 are activated by thrombin, whereas in mice PAR3 and PAR4 are the main receptor for thrombin. In addition, PAR2 is activated by trypsin, tryptase and other coagulation factors (35, 37–40). Thrombin-dependent activation of PAR1 occurs by proteolytic cleavage of the extracellular domain (35, 41, 42). In addition, other proteases including granzymes (A, B and K) activate PAR1 although the cleavage site is not well defined (42–49). Extracellular granzymes A, B and K have been shown to induce inflammation by activating PAR1 in multiple cell types including neurons, fibroblast and endothelial cells (43, 45–47, 49, 50).

While GZMs are viewed mainly involved in cytotoxic function, non-cytotoxic and extracellular activities have been reported including regulation of vascular integrity, angiogenesis, extracellular matrix remodeling (51–53). In contrast to thrombin, granzymes cleavage site and the downstream signaling pathways differ from thrombin leading to a non-canonical activation of PAR1 called biased signal (42–49, 54, 55).

In this study, we evaluated whether T_ang_ cells are involved in endothelial repair following acute lymphocytic choriomeningitis virus (LCMV) infection. We found that CD31^high^ expressing CD8 T_ang_ cells increased after viral control, have a memory precursor phenotype (KLRG1^-^CD127^+^) and secrete vascular endothelial growth factor (VEGF) suggesting their contribution to endothelial repair. Moreover, we found that in the setting of chronic HIV infection, immune activation drives a proinflammatory phenotype of CD8 T_ang_ that express granzymes. The frequencies of GZMA^+^GZMB^+^CD8 T_ang_ are associated with the atherosclerotic cardiovascular disease (ASCVD) risk. In addition, plasma levels of GZMA correlated with endothelial inflammatory markers including sVCAM1 and IL-6. *In vitro* GZMs activate PAR1 favoring the secretion of proinflammatory cytokines. This evidence supports the notion that HIV driven immune activation alters CD8 T_ang_ cell function and chronic stimulation of this pathway may contribute endothelial inflammation and the increased risk of cardiovascular disease.

## Material and Methods

### Study Participants

Participants were studied under a MedStar Georgetown University Hospital and NIAID Institutional Review Board approved HIV clinical research studies. Characteristics of the study groups of PWoH (n= 17) and PWH (n= 25) are described in Table 1. All study participants signed a written informed consent for the collection of samples. Animal studies were conducted under Georgetown University IACUC approved protocol.

**Table 1.** Study groups characteristics.

| **Demographics** | **PWH (n= 25)** | **PWOH (n= 17)** |
| --- | --- | --- |
| Age, yr, median (IQR) | 52 (42-57.5) | 48 (34-60) |
| Gender n (%) |  |  |
| Male | 22 (88) | 8 (47.1) |
| Female | 3 (12) | 9 (52.9) |
| Ethnicity n (%) |  |  |
| White | 8 (32) | 11 (64.8) |
| African American | 12 (48) | 3 (17.6) |
| Other | 5 (20) | 3 (17.6) |
| **Clinical Characteristics** |  |  |
| Smoker ever, n (%) | 8 (32) | N/A |
| Diabetes, n (%) | 9 (36) | N/A |
| ASCVD risk estimator, median (IQR)<sup>1</sup> | 7.9 (5.0-18.6) | N/A |
| **Clinical Laboratory, median (IQR)** |  |  |
| Cholesterol, mg/dL<sup>2</sup> | 168 (147-194) | N/A |
| HDL, mg/dL<sup>3</sup> | 47 (41-56.3) | N/A |
| LDL, mg/dL<sup>3</sup> | 99.5 (78.5-117.5) | N/A |
| **HIV Characteristics, median (IQR)** |  |  |
| Viral Load copies/mL<sup>4</sup> | <20 (20-20) | N/A |
| CD4 count, cells/μL | 663 (532-861.5) | N/A |
| **ART treatment** |  |  |
| NRTIs, n (%) | 19 (76) | N/A |
| NNRTIs, n (%) | 4 (16) | N/A |
| PIs, n (%) | 4 (16) | N/A |
| INSTIs, n (%) | 14 (56) | N/A |
| **Serum Biomarkers, median (IQR)** |  |  |
| sVCAM-1, pg/mL | 288.5 (188.6-366.6) |  |
| IL-6 | 3.7 (1.2-4.6) |  |
| Granzyme A | 256.1 (111.1-514) |  |
| Granzyme B | 920.8 (523.4-1566) |  |
| IFNγ | 289.8 (87.0-1012) | |
| TNFα | 2.5 (1.1-4.6) | |
| sFASL | 9.8 (3.6-24.1) |  |

<sup>1</sup> 5 out of 25 participants didn't have ASCVD risk estimator. <sup>2</sup> 10 out of 25 participants didn't have total cholesterol. <sup>3</sup> 11 out of 25 participants didn't have HDL and LDL. <sup>4</sup> 4 out of 25 participants have viral load 110, 70, 60 and 50 HIV RNA copies/mL respectively.

### C57BL/6 f2r^fl/fl^ Mice

The mouse carrying a conditional floxed (fl) PAR1 (C57BL/6 F2r^fl/fl^) was constructed with Ingenious Targeting Laboratories (NY, USA). Exon 2 of F2r gene was conditionally targeted and it is fused in frame with p2A-Tomato reporter cassette. F2r and tdTomato genes are inactivated upon Cre mediated deletion between the LoxP elements. A FRT/LoxP selection ivNeo marker as homology arms were included in the final targeting vector.

Targeted iTL IN2 (C57BL/6) embryonic stem cells were microinjected into Balb/c blastocysts. Resulting chimeras with a high percentage black coat color were mated to C57BL/6 FLP mice to generate Somatic Neo Deleted mice. Confirmed somatic neomycin cassette deleted mice were bred with wild-type C57BL/6N mice to generate germline neomycin cassette deleted floxed PAR1 mice C57BL/6 f2r^fl/fl^.

All mice were bred and maintained under specific pathogen-free conditions at Georgetown University Division of Comparative Medicine facility. All experimental procedures were conducted in accordance with the guidelines and an approved protocol by Institution Animal Care and Use Committee at Georgetown University. Mice between 8 to 12 weeks old were used for experiments.

### RT-qPCR analysis

qPCR was performed to determined mRNA expression of PARs in resting murine cells. RNA was obtained from magnetic isolated B, CD4 and CD8 T cells (Miltenyi Biotech, MD). RNA was extracted and purified using RNeasy Mini kit (Qiagen, MD). RNA was reverse transcribed into cDNA using QuantiNova Reverse Transcription kit (Qiagen, Germany). qPCR performed with QuantiNova SYBR Green PCR kit in CFX Opus Real-Time PCR system (Bio-Rad, CA). Primers used: murine *F2r* (NM_010169.3): forward, 5′-ATGAAAGCGTCCTGCTGGAG-3′; reverse, 5′- GGACGTTCAGAGGAAGGCTG-3′; murine *F2rl1* (NM_007974.4): forward, 5′- GCTGGGAGGTATCACCCTTCT-3′; reverse, 5′-CGCAGAGAACTCATCGATGGA-3′; murine *F2rl2* (NM_010170.4): forward, 5′-TGCCAGTCACTGTTTGCCAA-3′; reverse, 5′- CTCGGGACACTCCGCTTTTAT-3′; murine *F2rl3* (NM_007975.3): forward, 5′- GACCCCCAGCATCTACGATG-3′; reverse, 5′-CAGCAGCAGTGCTTGAGAGCT-3′. Housekeeping gene HPRT was used: murine HPRT (NM_013556): forward, 5′ - GCGATGATGAACCAGGTTATGA- 3′; reverse, 5′ - ACAATGTGATGGCCTCCCAT - 3′ LCMV Glycoprotein: forward, 5′- CATTCACCTGGACTTTGTCAGACTC -3′; reverse, 5′- GCAACTGCTGTGTTCCCGAAA -3′ (56, 57).

Ct of each sample obtained from the amplification curve was normalized to HPRT and calculated ΔCt (Ct gene of interest – Ct HPRT). Results were expressed as 2^-ΔCt^ value.

### LCMV infection

C57BL/6-F2r^fl/fl^ mice were intravenously infected with 2x10^5^ pfu LCMV Armstrong strain. At days 6 and 21 post infection (p.i.), mice were anesthetized and perfused with cold HBSS before organs (lung and spleen) were collected. *In vivo* CD8 T cell depletion was performed in LCMV infected mice at day 15 p.i. Mice were injected intraperitoneally with 300 μg of anti-mouse CD8α mAbs (YTS 169.4, BioCell, MA) or IgG2b isotype control (LTF-2, BioCell, MA). CD8 T cell depletion was confirmed by flow cytometry in lung and spleen.

Cell suspensions from the spleens were obtained by smashing the spleens with a syringe plunger in PBS and red blood cells were lysed with ACK (Gibco, MA) for 3 min on ice. After cell lysis cell suspensions were passed through a 70 μm mesh, then cell counts and viability were assessed.

Cell suspension from the lungs were obtained by enzymatic digestion. Lungs were cut into small pieces and digested using Lung Dissociation kit (Miltenyi Biotech, MD). Cell suspensions were passed through a 70 μm mesh, then cell counts and viability were assessed. Total cell suspensions were used to analyze endothelial cells in the lung. For T cell analysis immune cell enrichment was performed using a 35% Percoll gradient. Total lung cell suspensions were centrifuged and pellets were resuspended in 12 ml of 35% Percoll (Sigma, MA) and centrifuged for 15 min without a brake at 1800 rpm (757g) at 7 °C. Immune cell suspensions were passed through a 70 μm mesh, then cell counts and viability were assessed.

### Flow cytometry

*Phenotype lymph organs and endothelial cell phenotype*: 2x10^6^ cells from spleen, lymph nodes and lung cells (pre-Percoll) were stained with LIVE/DEAD (Invitrogen, CA) for 30 minutes at 4°C. Cells were incubated with a cocktail of mAbs (Table S1, panel-1 and panel-2) for 20 minutes at 4°C. Cell were washed with staining buffer (HBSS with 0.01% of NaN_3_ and 0.01% of BSA) twice and acquired with a BD Symphony (BD Biosciences, CA). Cells were analyzed with FlowJo (BD Biosciences, CA).

*Cytokine secretion assay*: 2x10^6^ splenic and immune lung cells (post-Percoll) were stimulated *in vitro* with 1 μg/mL of LCMVGp33-41 peptide (KAVYNFATC, Anaspec, CA) in the presence of 5 μg/mL of Brefeldin A (Sigma, MA) and 50 U/mL of rhIL-2 (National Cancer Institution, MD) for 4 hours at 37°C and 5% CO_2_. After stimulation, cells were stained with LIVE/DEAD (Invitrogen, CA) for 30 minutes at 4°C. Cells were washed twice and incubated with surface mAbs cocktail (Table S1, panel-3) for 20 min at 4°C. Cells were fixed and permeabilized with BD Cytofix/Cytoperm solution (BD Bioscience, CA) and then incubated with the mAbs cocktail for intracellular markers for 20 min at 4°C (Table S2 panel-2). Samples were acquired in a BD Symphony and analyzed with FlowJo (BD Biosciences, CA).

*MHC class I pentamer staining*: 1x10^6^ splenic and lung immune cells from infected mice were stained with H-2D^b^ restricted LCMVGP33-41aa pentamer (H-2D^b^-KAVYNFATC, Proimmune, UK) at room temperature (22 °C) for 10 min followed by incubation with surface mAbs cocktail (Table S1) for 20 min at 4°C. Cell suspensions were acquired in a BD Symphony (BD Biosciences, CA) and analyzed using FlowJo.

*CD8 T_ang_ sorting and cytokine secretion*: T cells were isolated from the lungs as described above. Lung cell suspensions from naïve and infected mice at day 21 p.i. were stained with a cocktail of mAbs for 20 minutes at 4°C (Table S1), and live cells were selected using Sytox Blue staining before sorting (Biolegend, CA). CD31^high^ and CD31^neg^ CD8 T cells were sorted on a FACSAria (BD Bioscience, CA). T cells were stimulated at a concentration of 300,000 cells/mL in plates with cross-linked anti-CD3 and anti-CD28 mAbs (BD Bioscience, CA). After 3 days of culture, supernatants were collected and cytokines (M-CSF, PDGF-A, GM-CSF, IL-2, IL-4, IL-5, IL-13, IL-17A, IL-6, IL-22, IL-9 and IL-10) were measured using LEGENDplex (Biolegend, CA) following manufacturer’s instructions. Briefly, 25 μL of sample in duplicates were incubated with capture mAbs coated beads for 2 hours followed by incubation with detection mAbs for 1 hour at room temperature in a shaker. Each sample was incubated with streptavidin-PE for 30 min and acquired with a BD Symphony (BD Biosciences, CA). Results were analyzed using cloud-based software Qognit (Qognit Inc, CA).

### Flow cytometry human peripheral blood mononuclear cells

Whole blood from people without HIV PWoH (n= 17) and PWH (n= 25) were collected in BD Vacutainer blood collection tube containing EDTA (BD Bioscience, CA). Whole blood was diluted with PBS with 2% FBS (1:1) and loaded on the top of Ficoll-Paque (Cytiva, MA) followed by a centrifugation at 400xg for 30 min at room temperature (25°C). Peripheral blood mononuclear cells (PBMCs) were collected and stained with LIVE/DEAD (Invitrogen, CA) for 30 minutes at 4°C. Cells were incubated with 1μg/mL human IgG (Sigma, MO) for 10 minutes at 4°C to block Fc receptors followed by incubation with a monoclonal antibodies (mAbs) cocktail (Table S2, panel-1) for 30 minutes at 4°C. Samples were acquired with a BD Symphony (BD Biosciences, CA) and analyzed using FlowJo software.

### Measurements of cytokines and biomarkers of endothelial inflammation in plasma

Plasma was obtained from whole blood collected in BD Vacutainer blood collection tubes containing EDTA (BD Bioscience, CA) from PWH (n= 25) and stored at -80°C until use. Cytokines, biomarkers of endothelial inflammation and granzymes (VCAM-1, IL-6, TNFα, IFNγ, sFasL, IL-2, granzyme A and granzyme B) were measured using LEGENDplex (Biolegend, CA) following manufacturer’s instructions as described above.

### Unsupervised clustering analysis

Multidimensional analysis was performed using Phenograph algorithm (FlowJo, CA). 10,000 of manually gated lung endothelial cells (CD326^-^CD45^-^CD31^+^) were concatenated from naïve mice and infected mice at days 6 and 21 p.i. Samples acquired on different days were normalized using Mutual Nearest Neighbors (MNN) (FlowJo, CA) (58). Unsupervised clusters were analyzed with Phenograph (k= 100) based on surface markers (CD31, PDL1, tdTomato, CD34, CD309, VCAM-1, CD105, CD90.2 and CD201). Mean fluorescence intensity of each marker for all the clusters was presented in heatmap and median frequency of each cluster from naïve and infected mice were expressed as bubble plots.

### Co-culture of primary human umbilical vein endothelial cells (HUVEC) and CD8 T_ang_ cells

Frozen PBMCs from PWoH (n= 10) and PWH (n= 10) were thawed and rested for 2 hours. Cells were incubated with 1μg/mL human IgG (Sigma, MO) for 10 minutes at 4°C to block Fc receptors followed by a cocktail of mAbs incubation for 30 minutes at 4°C (Table S2, panel-2). Live cells were selected using Sytox Blue staining before sorting (Biolegend, CA). CD8 T_ang_ cells were sorted on a FACSAria (BD, CA) and centrifuged at 1000 rpm for 30 minutes. Sorted CD8 T_ang_ cells were activated with anti-CD3 and anti-CD28 mAbs coated beads (Miltenyi Biotech, MD) for 3 days and expanded in the presence of 50 U/mL of recombinant human (rh) IL-2 (National Cancer Institution, MD) for 14 days. 5,000 HUVEC and 100,000 of CD8 T_ang_ cells were resuspended in endothelial cell growth basal media (Lonza, CA) and seeded in the 96 well plate in the presence or absence of anti-CD3/anti-CD28 mAbs coated beads (Miltenyi Biotech, MD) and incubated at 37°C and 5% CO_2_ overnight. As a control, HUVEC alone was stimulated with 20 ng/mL of rhIFNγ (PeproTech, NJ) and 0.5 ng/mL of rhTNFα (R&D System, MN) overnight. Cytokines in supernatant were measured using LEGENDplex (Biolegend, CA) as described above.

### Tube formation assay

Frozen PBMCs from PWH (n= 3) and PWoH (n= 2) were thawed and rested for 2 hours and sorted as described above. Cells were incubated with 1μg/mL human IgG (Sigma, MO) for 10 minutes at 4°C to block Fc receptors followed by a cocktail of mAbs incubation for 30 minutes at 4°C (Table S2, panel 2). Live cells were selected using Sytox Blue staining before sorting (Biolegend, CA). CD8 T_ang_ memory cells (CD45RA^-^CD27^+^) were sorted as described above. For HIV-Gag-specific CD8 T cells, PBMCs from PWH were stimulated with HIV_gag_ peptide pool (2 μg/mL, NIH AIDS Reagent Program) overnight and CD137^+^CD8 T cells were sorted. Sorted cells were washed twice with cold PBS and labeled with 1.5 μM of CSFE (Invitrogen, CA) for 8 min at 37°C followed by quenching with cold FBS for 5 min on ice. CFSE labeled sorted cells were rested overnight. 10 μL of basement membrane extract (Cultrex reduced growth factor, R&D System, MN) was applied in the inner well of 15 well μ-slide chambered coverslip (ibidi, WI) and incubated for 20 min at 37°C for gelation. HUVEC and CFSE labeled CD8 T_ang_ cells were resuspend in endothelial cell growth basal media (Lonza, CA) at 2:1 endothelial cell to T cell ratio and seeded on the top of gel in each well. Chambered coverslips (Ibidi, WI) were incubated in a stage-top cell incubator at 37°C and 5% CO_2_ and evaluated using Leica SP8 live cell imaging (Leica, IL) at 4, 6, 8, 10 and 12 hours at 10X magnification. Analysis was analyzed using the image analysis platform Wimasis (Wimasis GmbH, Germany).

### Calcium mobilization assay

60,000 HUVEC were cultured on a coverslip in endothelial cell growth media (Lonza, CA) for 8 hours and starved with endothelial cell growth basal media-2 (EBM-2) containing 1% PBS overnight. Cells were loaded for 40 min with 1 µM Fluo-4-AM (Invitrogen, Thermo Fisher Scientific) in a buffer solution containing 140 mM NaCl, 4 mM KCl, 1 mM MgCl_2_, 1.2 mM CaCl_2_, 10 mM HEPES, and 5 mM glucose (pH 7.3). Cells were imaged with 20X objective using a Nikon TE2000 microscope with an excitation filter of 480 ± 15 nm and an emission filter of 535 ± 25 nm. The images were captured by a Retiga 3000 digital camera (QImaging) and analysis was performed offline using ImageJ.

### Granzyme-dependent PAR1 activation

100,000 HUVEC were seeded a 12 well plate in endothelial cell growth media supplemented with EGM-2 bullet kit overnight. Next day cells were starved in EBM-2 containing 1% FBS for 6 hours. HUVEC were incubated pre-incubated with 10 μg/mL of ATAP-2 (Invitrogen, CA) or 300 nM of PAR1 antagonist SCH530348 diluted in EBM-2 containing 1% FBS for 20 min at room temperature (25°C). After the preincubation period, supernatants were removed from the wells, and HUVEC were stimulated with 100 nM of recombinant human granzyme A (rhGZMA, Kamiya Biomedical, WA) and recombinant human granzyme K (rhGZMK, Kamiya Biomedical, Japan) with fresh inhibitors at the same concentration. As the positive control, HUVEC were incubated with 100 μM of PAR1 agonist TFLLR-NH2 (Tocris Bioscience, UK) and control peptide RLLFT-NH2 (Tocris Bioscience, UK). After overnight incubation at 37°C and 5% CO_2_ supernatants were collected, and cytokines were measured using LEGENDplex (Biolegend, CA) as described above.

### Statistical analysis

Comparisons were performed using non-parametric Mann-Whitney test or Wilcoxon test and *p* value ≤ 0.05 was considered significant. Correlations were performed using nonparametric Spearman correlation and *p* value ≤ 0.01 was considered significant.

## Results

### PAR1 expression is modulated in virus-specific CD8 T lymphocytes during acute LCMV infection

At the site of vascular injury mediated by trauma or infection, the crosstalk between the coagulation cascade, platelets, endothelial cells and CD8 T cells is mediated by thrombin dependent activation of PAR1 (30–32, 35). Particularly, PAR1 signaling is involved in CD8 T cell trafficking and function (30, 33). However, whether PAR1 is modulated in CD8 T cells during the immune response remains to be defined.

We generated a PAR1 reporter mouse that allows us to evaluate PAR1 expression in CD8 T cells during the course of a systemic viral infection and better understand the interplay between CD8 T cells and endothelial cells. C57BL/6 F2r^fl/fl^ mouse carrying floxed Exon 2 of *F2r* gene was conditionally targeted as well as fused in frame with p2A-Tomato (tdT) reporter cassette (as described in Material and Methods section and Figure S1A).

C57BL/6 F2r^fl/fl^ mice showed similar cell numbers in lymph nodes and spleen compared with the wild type C57BL/6 (Figure S1B). The frequencies and cell count of innate cells including neutrophils, eosinophils, monocytes, macrophages and dendritic cell subsets were similar between the C57BL/6 F2r^fl/fl^ and the C57BL/6 wild type mice (Figure S1C and D). Similarly, no significant differences between C57BL/6 F2r^fl/fl^ and wild type mice were observed in the frequencies and cell numbers of lymphocytes subsets, B, NK and T cells in lymphoid organs (Figure S1E).

We next examine PAR1 expression by determining the frequency of tdTomato^+^ (tdT^+^) cells, immune cells. TdTomato was found in CD4 and CD8 T cells and NK cells but not in B cells (Figure 1A, B and C). Similar to our previous observations in human T cells, CD8 T cells expressed higher levels of PAR1 compared with CD4 T cells. Among lymphocytes, NK expressed the highest level of PAR1 under steady state conditions (Figure 1C) (33).

**Figure 1.**
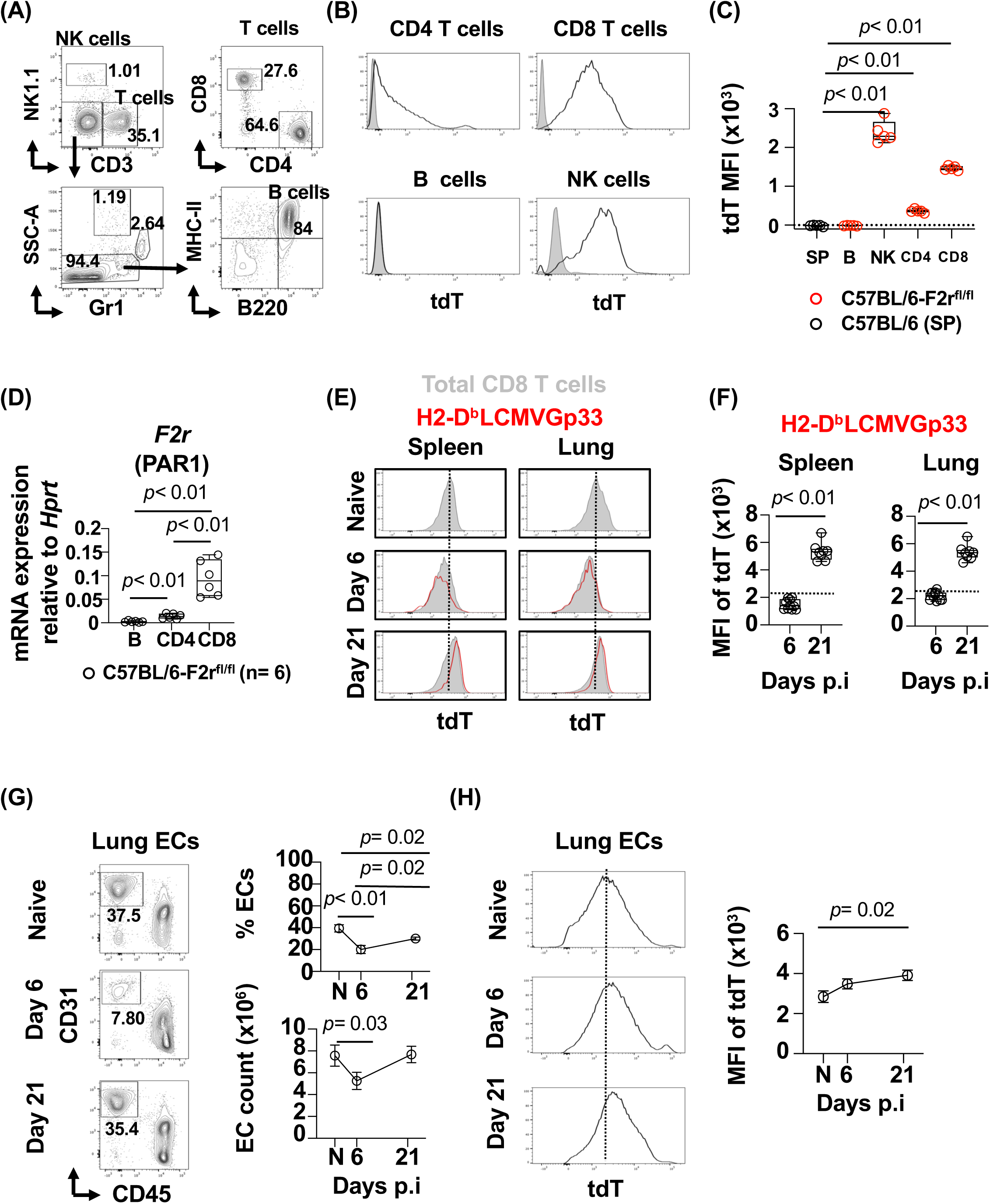
PAR1 expression is modulated in LCMVGp33-specific CD8 T lymphocytes during acute LCMV infection. *In vivo* expression of PAR1 in splenic cells from naïve C57BL/6-F2r^fl/fl^ (n= 5) and C57BL/6 (n= 5) mice analyzed by flow cytometry. **(A)** Cells were analyzed after gating on live cells, and singlets and CD45^+^ lymphocytes. Representative contour plots gating cell types: NK cells (CD3^-^NK1.1^+^), B cells (MHCII^+^B220^+^), CD4 (CD3^+^CD4^+^) and CD8 T cells (CD3^+^CD8^+^). **(B)** Representative histogram of tdTomato (PAR1) expression in CD4 and CD8 T cells, B cells and NK cells from C57BL/6 mice (solid grey) and C57BL/6-F2r^fl/fl^ mice (black line). **(C)** Median Fluorescence Intensity (MFI) of tdTomato expression of total splenic cells from C57BL/6 mice (SP) and B cells, NK and CD4 and CD8 T cells and NK cells from C57BL/6-F2r^fl/fl^ mice. **(D)** Relative *F2r* mRNA expression in sorted B cells and purified by magnetic isolation of CD4 and CD8 T cells from C57BL/6-F2r^fl/fl^ mice (n= 6). **(E)** C57BL/6-F2r^fl/fl^ mice were infected with LCMVArm strain (2 x 10^5^ pfu, i.v). Splenic and lung cells were assessed at day 6 and 21 p.i. by flow cytometry. **(F)** Representative histogram of tdTomato (tdT) expression in total CD8 T cells from naïve (N) and infected mice at day 6 and 21 p.i. (solid gray). Overlayed, tdTomato expression in LCMVGp33-specific CD8 T cells detected by H2-D^b^LCMVGp33 pentamer staining (red line). **(F)** Median Fluorescence Intensity (MFI) of tdTomato expression in LCMVGp33-specific CD8 T cells detected by H2-D^b^LCMVGp33 pentamer staining. Dashed line represents level expression of total naïve CD8 T cells. **(G)** Representative plot of lung endothelial cells (EC). After excluding dead cells and doublets, ECs were gated as CD326^-^CD45^-^CD31^+^. Frequency and cell count of lung endothelial cells from naïve (N) and infected mice at days 6 and 21 p.i. Frequencies of ECs are expressed as frequency of total live cells. **(H)** Median Fluorescence Intensity (MFI) of tdTomato expression in lung ECs from naïve (N) and infected mice (day 6 and day 21p.i.). The bar graph is represented by box and whisker showing the median value with first and third quartiles in the box, with whiskers extending to the minimum and maximum values. The data in line graphs represent mean±SEM in each time point. Data are pooled from three independent experiments with 5-9 naïve animals (C, E and F) and 8-17 per time point (F and H). Statistical analysis was performed using non-parametric Mann-Whitney test. *P* value < 0.05 was considered significant.

mRNA of *F2r* expression in tdT expressing cells was further confirmed by qPCR in magnetic cell isolated CD4 T cells, CD8 T cells and B cells. CD8 T cells expressed higher levels of mRNA of *F2r* than CD4 T cells (Figure 1D). PAR1 protein expression in CD4 and CD8 T cells was further confirmed by flow cytometry in tdT^+^ T cells (Figure S2). These data show that under steady state conditions, PAR1 is expressed in T and NK cells but not B cells.

We next investigated the expression of PAR1 in virus-specific CD8 T cells during a systemic acute viral infection with the lymphocytic choriomeningitis virus Armstrong strain (LCMVArm). This infection model recapitulates features of human viral infections, the acute form, LCMV Armstrong (Arm) is cleared acutely at 8-10 days post infection (p.i) and induces the generation of LCMV-specific memory CD8 T cells (59–61). LCMV is a non-cytolytic virus therefore the pathology observed in infected hosts is mediated by the host immune response (62, 63).

Using this model, we aimed to study the inflammation mediated endothelial injury in the lung, and address whether T cells are involved in endothelial repair after viral control. We studied two time points post-infection: day 6 p.i (effector phase) and day 21 p.i (contraction phase and early memory development) (61, 64). LCMVGp33-specific CD8 T cells secreted IFNγ, TNFα and showed cytotoxic potential (as measured by CD107a surface expression) upon *in vitro* stimulation with LCMVGp33-41aa peptide (Figure S3A and B). LCMVGp33-specific CD8 T cells peaked by day 6 p.i in both the spleen and lung, then declined by day 21 in the spleen when the virus was controlled (Figure S3A and B). In contrast, LCMVGp33-specific CD8 T cells in the lung, remained elevated (Figure S3A and B).

Expression of PAR1 was monitored by tdT^+^ cells in total CD8 T cells and virus-specific CD8 T cells using pentamer staining (H2-D^b^LCMVGp33-41aa) (Figure S3C and Figure 1E respectively). At day 6 p.i. and compared to total CD8 T cells from the naïve (dashed line) mice, expression of PAR1 (tdT^+^) in LCMVGp33-specific CD8 T cells was reduced, but was significantly upregulated by day 21 p.i. (Figure 1E and F). Similar observations were noted in total CD8 T cells in both spleen and lung (Figure S3C and 1E). PAR1 transcriptional regulation in LCMVGp33-specific CD8 T cells may be induced by *in vivo* granzyme activation during degranulation (Figure S3B) (33). Consistent with previous observations in degranulating human CD8 T cells, tdT expression by CD107a^+^CD8 T cells showed lower expression of granzyme A compared to CD107a^-^CD8 T cells (Figure S3E) (33). In addition, tdT^+^CD8 T cells negatively correlated with the frequency of GZMA^+^CD8 T cells, R= -0.61, *p*= 0.02.

Altogether these data show that PAR1 is regulated at distinct stages of CD8 T cell activation, with lower PAR1 expression at the effector phase and upregulated after viral control and memory development.

### Angiogenic CD31^high^CD8 T (T_ang_) cells promote endothelial recovery during acute viral infection

In humans, T_ang_ cells were originally described in the center of endothelial progenitor cell colonies isolated from peripheral mononuclear cells (PBMCs). In *in vivo* experiments, T_ang_ cells support new vessel formation (25). However, whether T_ang_ cells are virus-specific CD8 T cells that acquire these properties once the infection is controlled and are involved in endothelial repair remains poorly understood. We hypothesized that systemic LCMVArm infection and its associated inflammation induces endothelial injury in the lung, and T_ang_ cells play a role in repair after viral control. We found at day 6 p.i., when LCMV-specific CD8 T cells are recruited into the lung, there was a significant reduction in the frequency and cell count of lung endothelial cells (ECs) (Figure 1G). ECs recovered to baseline levels at day 21 p.i. (Figure 1G). In contrast to CD8 T cells, expression of PAR1 in endothelial cells increased over the course of infection suggesting that PAR1 can be targeted by different proteases during the course of the infection (Figure 1H). These data show that a mild endothelial cell injury occurs in lung at the peak of CD8 T cell response and recovers upon viral control.

We next determined if CD8 T cells are involved in the recovery of lung ECs. LCMV infected mice were either treated with anti-CD8 mAb and IgG control at day 15 p.i. when the infection is controlled. CD8 T cells were efficiently depleted in the lung by day 21 p.i. (Figure 2A). Compared to the mice treated with IgG control, CD8 T cell depleted mice showed a delayed recovery of EC count suggesting that CD8 T cells are involved in this process (Figure 2B). At this time point of infection, we did not detect LCMV glycoprotein by qPCR in both spleen and lung suggesting viral control (Figure S4A) (65).

**Figure 2.**
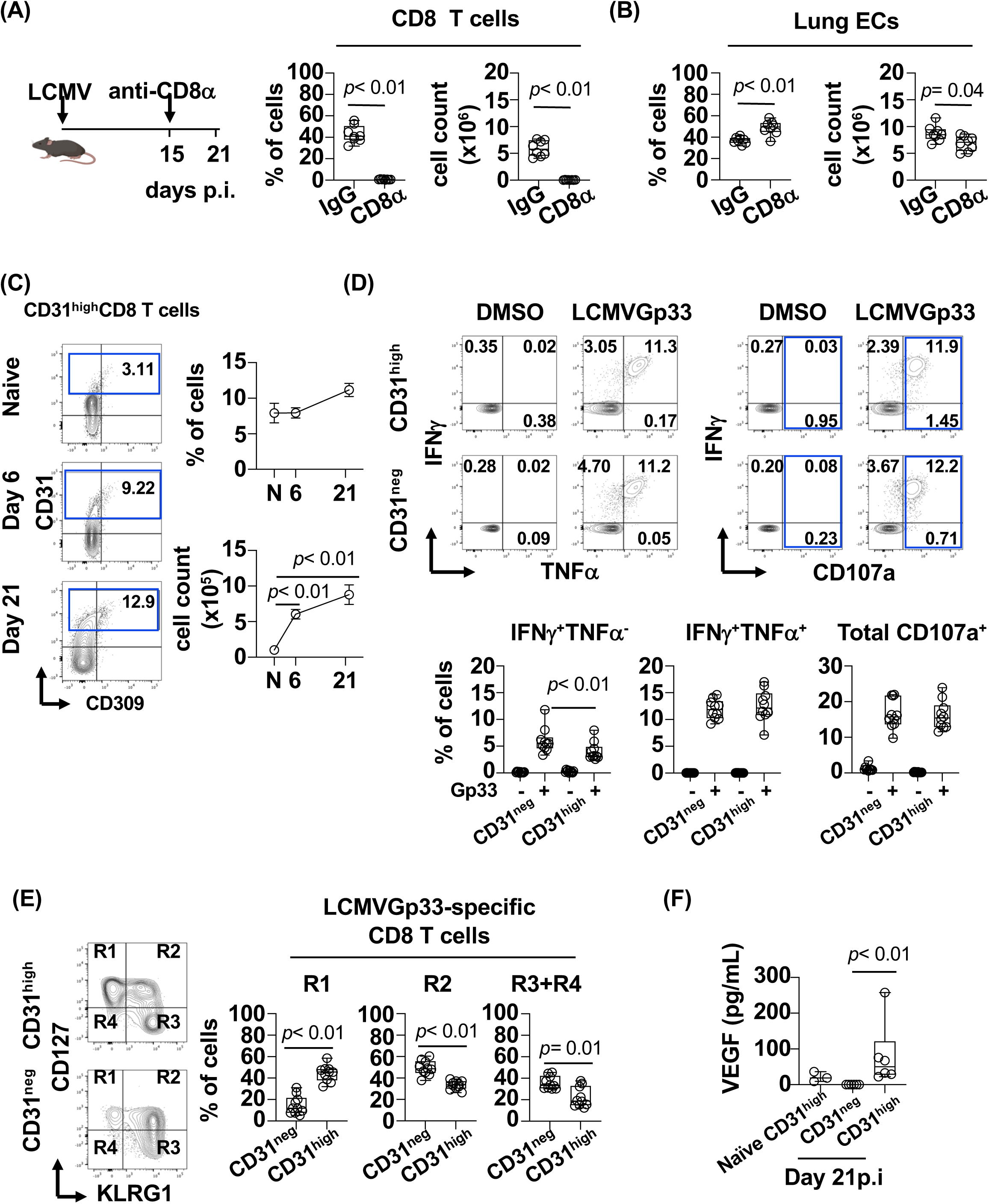
Angiogenic CD8 T cells promote endothelial recovery during acute viral infection. **(A)** Schematic presentation of CD8 T cell depletion. C57BL/6-F2r^fl/fl^ mice were infected with LCMVArm strain (2 x 10^5^ pfu, i.v). At day 15 p.i mice were administered anti-CD8α mAb and IgG2b isotype for control mice. Frequency and cell count of CD8 T cells at day 21p.i in CD8 T cell depleted mice and IgG treated control mice assessed by flow cytometry. **(B)** Lung endothelial cell frequency and count from CD8 T cell depleted and control mice at day 21 p.i. Frequencies of ECs are expressed as frequency of total live cells. **(C)** Representative contour plot of CD31^high^ and CD309 expression by lung CD8 T cells from naïve mice and infected mice. Frequency and cell count of CD31^high^CD8 T cells from naïve (N) mice and infected mice at day 6 and 21 p.i. **(D)** Lung CD31^high^ and CD31^neg^ CD8 T cells, representative contour plot of secretion of IFNγ, TNFα secretion and CD107a upon *in vitro* stimulation with LCMVGp33-44 peptide. DMSO was used as control. **(E)** Representative contour plot of KLRG1 and CD127 expression by LCMVGp33-specific CD8 T cells. Frequency of KLRG1^-^CD127^+^ (R1), KLRG1^+^CD127^+^ (R2) and KLRG1^+^CD127^-^ and KLRG1^-^CD127^-^ (R3+R4) LCMVGp33 specific CD8 T cells from lung at days 6 and 21 p.i.. **(F)** Lung CD31^high^ and CD31^neg^ bulk CD8 T cells were sorted from naïve mice and infected mice at 21 p.i. and stimulated *in vitro* with anti-CD3 and anti-CD28 mAbs coated plate for 3 days. Detection of VEGF in the supernatants of the culture. The bar graphs are represented by box and whisker showing the median value with first and third quartiles in the box, with whiskers extending to the minimum and maximum values. The data in line graphs represent mean±SEM in each time point. Data are pooled from two independent experiments with 8 naïve animals (A and B) and 7-16 per time point (C, D and E). Sorted cells from 3 naïve and 6 infected mice per time point (G). Statistical analysis was performed using non-parametric Mann-Whitney test. *P* value < 0.05 was considered significant.

We next tested the hypothesis that the delayed recovery of lung ECs is associated with the loss of CD8 T_ang_ cells during depletion. T_ang_ cells have been defined by expression of CD31 (25). In the lung, during the course of the LCMV infection, a small subset of CD8 T cells expressed CD31^high^ and CD309 (vascular endothelial growth factor receptor 2 (VEGFR2)) and although the frequency showed an increased trend by day 21 p.i., it did not rich statistical significance (Figure 2C). In contrast, the number of CD31^high^CD8 T cells was significantly increased at day 6 and 21 demonstrating an expansion of this subset during the recovery phase of lung ECs (Figure 2C). In addition, similar PAR1 expression was observed in both CD31^high^ and CD31^low^ CD8 T cells (Figure S4B).

We then investigated whether CD31^high^CD8 T cells were LCMV-specific and secrete cytokines upon *in vitro* stimulation with LCMVGp33 peptide (Figure 2D). At day 21 p.i. and compared to CD31^neg^CD8 T cells, CD31^high^CD8 T_ang_ cells secrete similar levels of IFNγ, TNFα and have similar cytotoxic potential (CD107a). Of note, the frequency of CD31^high^LCMVGp33-specific CD8 T cells single IFNγ producers were reduced compared to the CD31^neg^ (Figure 3D).

**Figure 3.**
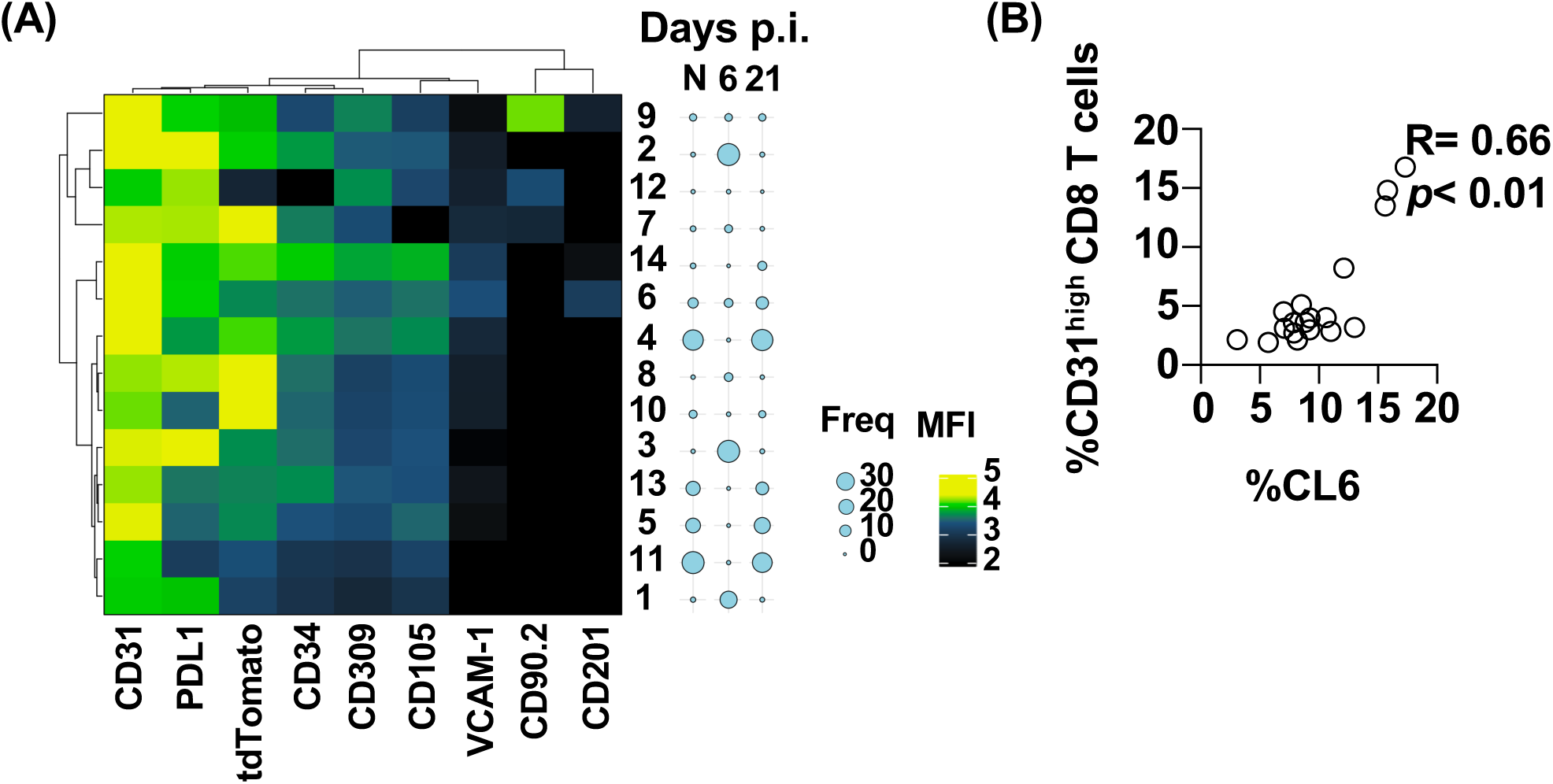
CD31^high^CD8 T cells are associated with endothelial that express a regenerative phenotype. **(A)** Unsupervised clustering analysis of lung endothelial cells using phonograph (k= 100). The heatmap represent mean fluorescence intensity of CD31, PDL1, tdTomato, CD34, CD309, VCAM-1, CD105, CD90.2 and CD201 expression in ECs. Bubble plots represent the mean frequency of each cluster from naïve and infected mice at day 6 and 21 p.i. **(B)** Relationship between frequency of Cluster 6 and lung CD31^high^CD8 T cells. Data are pooled from two independent experiments with 7 naïve (A and B) and 8-10 per time point animals (A and B). Correlation were performed using non-parametric Spearman correlation and *p* value ≤ 0.01 was considered significant.

Interestingly, LCMVGp33-specific CD31^high^CD8 T cells were enriched in the memory precursor effector T cells (KLRG1^-^CD127^+^), which have survival advantage during the contraction phase and possesses plasticity to give rise to multiple memory subsets (Figure 2E) (66–69). As expected, at day 21 p.i., a reduction of double positive effectors (KLRG1^+^CD127^+^) and short-live effector KLRG1^+^CD127^-^ and KLRG1^-^CD127^-^ CD8 T cells was observed at day 21 p.i. (Figure 2E).

In addition, we evaluated the ability of lung CD31^high^CD8 T cells to secrete pro-angiogenic cytokines and T cell helper cytokines. We sorted CD31^high^ and CD31^neg^ bulk CD8 T cells from lung of LCMV mice infected at day 21 p.i., and CD31^high^ bulk CD8 T cells from lung of naïve mice as control. Sorted subsets were TCR stimulated with cross-linked CD3 and CD28 mAbs for 3 days and cytokines were measured in the supernatants (Figure 2F and Figure S4C). CD31^high^ but not CD31^neg^CD8 T cells secreted vascular endothelial growth factor (VEGF), a cytokine that induces proliferation of ECs suggesting a pro-angiogenic function (70, 71). In addition, both CD31^neg^ and CD31^high^CD8 T cells showed similar ability to secrete granulocyte-macrophage-colony stimulating factor (GM-CSF), known to induce EC proliferation and mobilize endothelial progenitor cells from the bone marrow (Figure S4C) (72, 73). Lastly, CD31^high^CD8 T cells showed the ability to secrete of IL-2 and secrete low levels of Th2, Th17, Th22 and Th9 associated cytokines (Figure S4C).

These data suggest that angiogenic CD31^high^CD8 T_ang_ cells are virus specific with an enriched memory precursor phenotype. CD31^high^CD8 T_ang_ secrete VEGF suggesting may contribute with lung EC recovery after viral control.

### Endothelial recovery is associated to CD31^high^CD8 T_ang_ cells

We next investigated the relationships between CD8 T_ang_ cells and lung ECs during the course of the LCMV infection. ECs are heterogenous and play important roles by supporting immune cell trafficking and regulating inflammation during immunity (74–76). Diverse phenotypes with distinct functions including “immune cell-like” and “developmental/regenerative” involved in immunity and repair respectively have been described (77, 78).

We hypothesized that during the phase of CD31^high^CD8 T_ang_ cells expansion and EC recover, ECs would be enriched with a regenerative phenotype. We determined the expression of markers associated with inflammation (PD-L1, VCAM1) and proangiogenic/repair (CD34, CD309, CD105) (63, 77, 79, 80). CD309 is a receptor for the VEGF, an essential factor for EC function during angiogenesis (70). CD105 (Endoglin) binds to different components of the TGF-β superfamily and also regulates endothelial proliferation, migration, and vascular homeostasis (81, 82). In addition, CD90.2 was used as maker for lymphatic endothelial cells (83).

We performed an unsupervised cluster analysis using PhenoGraph to classify high-parameter single-cell data into phenotypically distinct clusters (84). We concatenated 10,000 events per mouse of CD326^-^CD45^-^CD31^+^ lung ECs from naïve and LCMV infected mice from day 6 and 21 p.i. to analyze changes in endothelial cells (Figure 3A).

We found that the frequency of lung EC clusters (CL) 1, 2, 3, and small CL 7, and 8 significantly increased at day 6.p.i (Figure 3A, bubble plots and Figure S5). These CLs expressed higher levels of PD-L1 and distinct levels of CD31 and PAR1 (Figure 3A). Expression of PD-L1 suggest that endothelial cells interact with immune cells regulating their function and potential immune mediated pathology through PD1-PD-L1 interaction (85).

The CL4, 5, 6, and 13 were present in the naïve mice and at day 21 p.i. (Figure 3A and Figure S5). These clusters expressed markers associated with EC proliferation including CD105 (Endoglin), and CD309 (VEGFR2), A small EC subset (CL14), expressed high levels of CD105 and was expanded by day 21 p.i. suggesting a “regenerative” phenotype (Figure S5). In addition, CL10 and CL11 remained significantly reduced compared to the naïve mice. CL9 correspond to CD90.2^+^ lymphatic endothelial cells and its frequency remained constant through the course of the infection.

These data suggested that ECs are heterogenous and dynamic during a viral infection. The ECs in CL6 expressed CD201 (endothelial protein C receptor, EPCR), a marker associated with resident vascular endothelial cell progenitors and activated protein C (APC) dependent activation of PAR1 during angiogenesis (78, 86, 87). Importantly, CL6 expressed higher levels of CD105, CD309 and CD34 suggesting a proliferative and regenerative phenotype (79). Moreover, CL6 was positively associated (R= 0.66, *p*< 0.01) with the frequency of CD31^high^CD8 T_ang_ cells suggesting a potential interaction during vascular remodeling (Figure 3B).

Altogether these data suggest that CD31^high^CD8 T_ang_ cells may play a role in endothelial recovery by secreting VEGF promoting proliferation of endothelial cells.

### Circulating angiogenic CD8 T (T_ang_) cells from PWH have proinflammatory phenotype

In humans with chronic inflammatory diseases reduced frequencies of T_ang_ cells are associated with cardiovascular risk (25, 28, 29). The role of T_ang_ cells in the setting of human chronic viral infections such as HIV remains to be defined. Previously, we reported that circulating T_ang_ cells from PWH express the vascular homing receptor CX3CR1 suggesting their recruitment to the sites of endothelial inflammation (88). Given that T cell immune activation and systemic inflammation are the main features of chronic HIV infection, we hypothesized that this environment alters T_ang_ cell function promoting a proinflammatory function that contributes to endothelial inflammation. To address this question, we investigated circulating T_ang_ cells in PBMCs from PWH (n= 25) and uninfected controls people without HIV (PWoH, n=17). PWH had a median CD4 count 663 (532-861.5) cells/μL and HIV RNA viral load < 20 copies/mL (n= 20) and 50-110 copies/mL (n= 4). Study participants had a median ASCVD (Atherosclerosis Cardiovascular Disease) risk score of 7.9 (IQR: 5.0-18.6) (Table 1).

In humans, T_ang_ cells are defined as surface expression of CD31/PECAM-1 (Platelet Endothelial Cell Adhesion Molecule) and the chemokine receptor CXCR4 (receptor for the Stromal Cell Derived Factor-1) (*25, 89*). Flow cytometric analysis showed that in PWH there is a significantly reduction in the frequencies of circulating CD8 T_ang_ cells (*p*< 0.001). A trend, although not statistically significant was observed in CD4 T_ang_ cells as well (Figure 4A and B).

**Figure 4.**
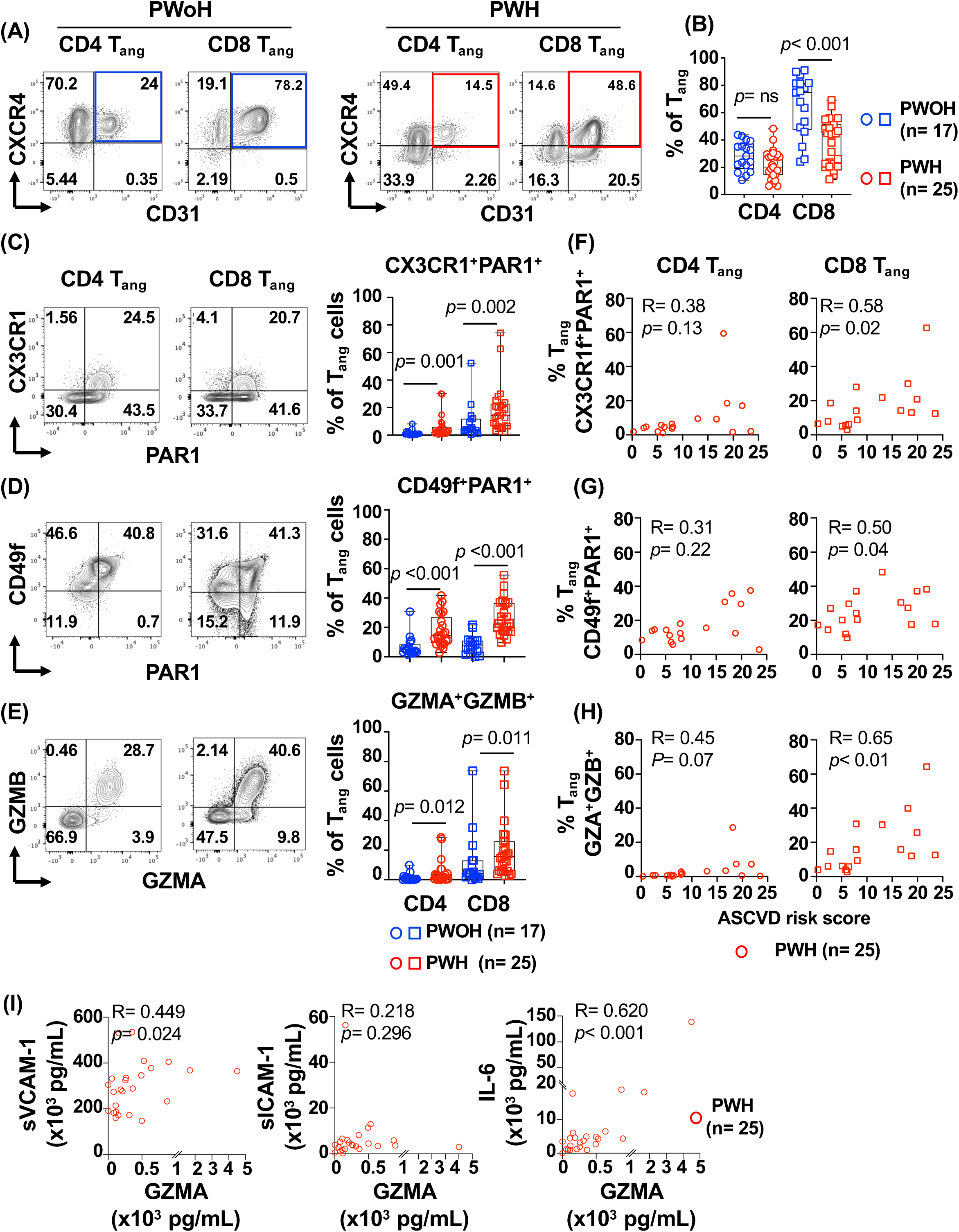
Proinflammatory phenotype of CD8 T_ang_ correlates to ASCVD risk in PWH. Circulating T_ang_ were analyzed in PBMCs from PWH (n= 25, red symbols) and PWoH (n= 17, blue symbols) by flowcytometry. After gating on live cells and singlets, CD4 and CD8 T cells were analyzed based on expression of CD31 and CXCR4: **(A)** Representative contour plots for gating angiogenic T cells, CD4 T_ang_ (CD31^+^CXCR4^+^) and CD8 T_ang_ (CD31^+^CXCR4^+^). **(B)** Frequency of CD4 T_ang_ and CD8 T_ang_ from PWH and PWoH. CD4 T_ang_ and CD8 T_ang_ expression of: (**C)** CX3CR1 and PAR1 **(D)** CD49f and PAR1 **(E)** granzyme A (GZMA) and B (GZMB). **F, G and H.** Relationship between expression of CD4 and CD8 T_ang_ cells and atherosclerotic cardiovascular disease (ASCVD) risk score in PWH. **(I)** Relationship between expression of plasma levels of sVCAM-1, sICAM-1, IL-6 and granzyme A in PWH (n= 25). The bar graph is represented by box and whisker showing the median value with first and third quartiles in the box, with whiskers extending to the minimum and maximum values. Statistical analysis was performed using non-parametric Mann-Whitney test. *P* value < 0.05 was considered significant. Correlations were performed using non-parametric Spearman correlation and *p* value ≤ 0.01 was considered significant.

To evaluate the impact of HIV driven immune activation, we studied expression of markers associated with adhesion, homing and trafficking during vascular inflammation including the PAR1 (30, 31, 33). We investigated the vascular homing receptor CX3CR1 that binds to CX3CL1 (fractalkine) expressed by inflamed endothelial cells, and the integrin CD49f (Integrin alpha-6) that binds to laminin, a component of the vascular basement membrane (30–32, 90). In PWH, both CD4 T_ang_ and CD8 T_ang_ cells showed significantly increased expression of PAR1, CX3CR1 and CD49f compared to PWoH suggesting an increased potential for recruitment and trafficking to the sites of vascular inflammation (Figure 4C and D). In addition to the increased trafficking potential, both CD4 T_ang_ and CD8 T_ang_ cells from PWH showed an increased expression of effector molecules Granzyme A (GZMA) and Granzyme B (GZMB) when compared to PWoH (Figure 4E).

All together these data suggest that PWH and virally suppressed by ART have a significant reduction in the frequencies of CD8 but not CD4 T_ang_ cells. T_ang_ cells showed a differentiated phenotype with cytotoxic potential and increased potential to migrate to the sites of vascular inflammation.

### Frequencies of activated CD8 T_ang_ cells are associated with cardiovascular risk in PWH

In PWH, immune activation, systemic inflammation and endothelial dysfunction have been identified as the main drivers of cardiovascular events (3, 5, 6, 16). The data above suggest that in the context of HIV infection T_ang_ cells have a proinflammatory phenotype. We next investigated the relationship between activation markers expressed by T_ang_ cells and the Atherosclerotic Cardiovascular Disease (ASCVD) risk score (Figure 4F, G and H).

The frequencies of circulating total CD4 and CD8 T_ang_ cells did not show an association with ASCVD. In contrast, we observed a positive correlation between ASCVD and the frequency of cells expressing activation markers including homing and trafficking CX3CR1^+^PAR1^+^CD8 T_ang_ and CD49f^+^PAR1^+^CD8 T_ang_ (R= 0.50, *p*= 0.04 and R= 0.58, *p*= 0.02 respectively) (Figure 4F and G). Similarly, GZMA^+^GZMB^+^CD8 T_ang_ cells showed a positive association with the ASCVD risk score R= 0.65, *p*< 0.01 (Figure 4H). Of note is that expression of these markers in CD4 T_angs_ cells did not correlate with ASCVD risk (Figure 4F, G and H). These observations suggest that in PWH immune activation of CD8 T_ang_ cells drives a proinflammatory phenotype and may play a role in sustaining endothelial inflammation.

To better address the potential contribution of activated CD8 T_ang_ cells on endothelial inflammation, we further studied the relationship between plasma levels of biomarkers of endothelial inflammation and immune cell trafficking including soluble Vascular and Intracellular Adhesion Molecule-1 (sVCAM-1 and sICAM-1 respectively), IL-6; and T cell-derived cytokines and effector molecules including TNFα, IFNγ, IL-2, sFASL, GZMA and GZMB (Figure 4I and Figure S6A). We found that plasma levels of GZMA but not GZMB were positively associated with IL-6 (R= 0.62, p< 0.001). sVCAM-1 showed a positive trend (R= 0.449, *p*= 0.024) but not with sICAM-1 (Figure 4I). As expected, plasma levels of cytokines (TNFα, IFNγ, sFASL, IL-2) positively correlated with the plasma levels of granzymes (Figure S6A).

Altogether the data shows that chronic HIV driven immune activation increase trafficking and proinflammatory properties of CD8 T_ang_ cells. Moreover, the association with ASCVD suggest that CD8 T_ang_ cells may contribute to endothelial inflammation, dysfunction and disease risk.

### *In vitro,* CD8 T_ang_ cells from PWH showed enhanced ability to induced sICAM-1 and VCAM-1 by endothelial cells

The above results suggest that chronic HIV infection alters CD8 T_ang_ cells skewing their function to pro-inflammatory. Because activated CD8 but not CD4 T_ang_ cells showed association with cardiovascular risk, in the following sections of the manuscript we focused our study on CD8 T_ang_ cells. To address the potential mechanisms involved by which CD8 T_ang_ cells drive endothelial inflammation, we first determined the effects of *in vitro* activated CD8 T_ang_ cells on primary human umbilical vein endothelial cells (HUVEC). In the supernatants of the co-cultures, we measured adhesion molecules including sICAM-1 and sVCAM-1, proinflammatory cytokines IL-6, angiopoetin-2 (Ang-2) and IL-8 (CXCL8); and pro-angiogenic cytokines, basic Fibroblast Growth Factor (bFGF) and Vascular Endothelial Growth Factors (VEGF) (91–93).

CD8 T_ang_ cells from PWH (n= 10) and PWoH (n= 10) were sorted based on the expression of CD31, CXCR4 and memory phenotype. CD8 T_ang_ cells were *in vitro* expanded with anti-CD3/CD28 mAbs and rhIL-2 (50 U/ml) for 14 days. Expanded CD8 T_ang_ cells were co-cultured with HUVEC at 20:1 CD8 T_ang_ to HUVEC ratio and stimulated in media (control) and with anti-CD3/CD28 mAbs coated beads overnight. CD8 T_ang_ co-cultured with HUVEC cells in media alone showed negligible levels of shedding of sICAM-1 and sVCAM-1. In contrast, when CD8 T_ang_ cells were TCR stimulated a significant increase of sICAM-1 and sVCAM-1 was detected in the supernatant of the cultures (Figure 5A). The effects mediated by CD8 T_ang_ cells from PWH were significantly higher than those observed with CD8 T_ang_ cells from PWoH (Figure 5A). Moreover, stimulation of CD8 T_ang_ from PWH led to an increased trend, although did not reach statistical significance, in the production of proinflammatory and pro-angiogenic cytokines IL-6, and bFGF respectively. No differences were observed in the secretion of IL-8, Ang-2 or VEGF. (Figure 5B). The combination of cytokines IFNγ (10 ng/mL) and TNFα (0.5 ng/mL) used as positive control in HUVEC cultured alone, showed no statistically significant effect on the shedding of sVCAM-1, sICAM-1. In addition, low IL-6 secretion was observed suggesting that CD8 T_ang_ cell-HUVEC contact dependent mechanism is involved in mediating shedding of adhesion molecules and IL-6 secretion by endothelial cells (Figure S6B and Figure 5B). Similar levels of Ang-2, IL-8/CXCL8 and the pro-angiogenic cytokines VEGF and FGFb were detected when endothelial cells were stimulated with combination IFNγ and TNFα (Figure S6B).

**Figure 5.**
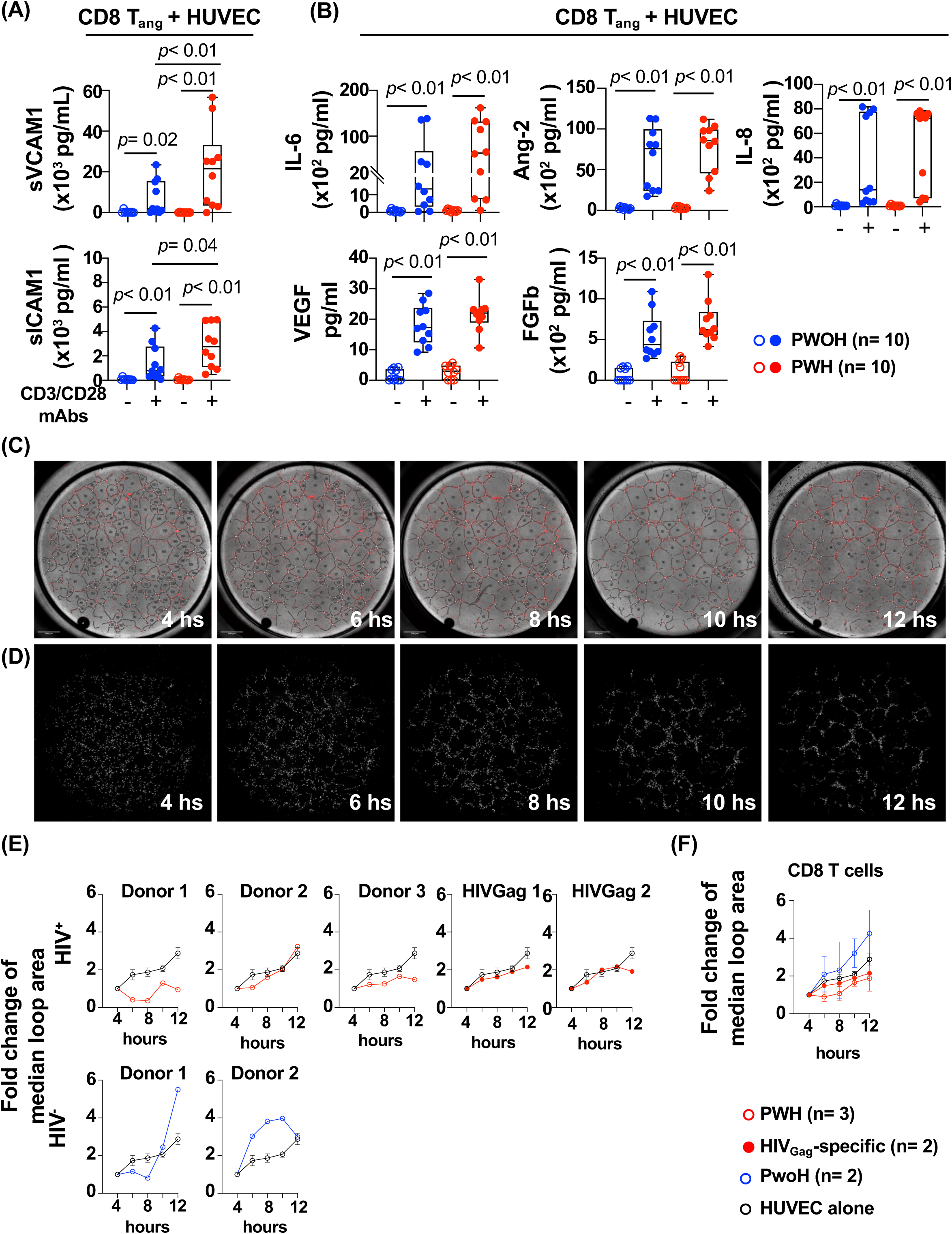
*In vitro*, CD8 T_ang_ cells from PWH induced shedding of sICAM-1 and sVCAM-1 by endothelial cells. CD8 T_ang_ from PWH (n= 10, red symbols) and PWoH (n= 10, blue symbols) were sorted and stimulated with CD3 and CD28 mAbs coated beads for 3 days and expanded in the presence of IL-2 for an additional 14 days. Activated CD8 T_ang_ cells were co-cultured with primary human umbilical vein endothelial cells HUVEC at 2 to 1 CD8 T_ang_ to HUVEC ratio for 24 hours. Supernatants were collected and measured: **(A)** Soluble vascular cell adhesion molecule 1 (sVCAM1) and intracellular adhesion molecule 1 (sICAM1). **(B)** Cytokines IL-6, Angiopoietin-2 (Ang-2), fibroblast growth factor (bFGF), vascular endothelial growth factor (VEGF) and IL-8/CXCL8. **(C)** Sorted CD8 T_ang_ memory cells (CD45RA^-^CD27^+^) from PWH (n= 3, opened red symbols), PWoH (n= 2, blue symbols), and HIVGag-specific CD8 T cells from PWH (n= 2, solid red symbols) were labeled with CFSE and cultured with HUVEC at a 2 to 1 ratio in Ibidi chambers for up to 12 hours. As control, HUVEC were cultured alone (black symbol). Representative phase contrast images at 2, 4, 6, 8, 10 and 12 hours with masks of tube (black line), branching point (white point), loop (labeled with number) and CD8 T_ang_ cells (red dots). **(D)** Representative fluorescent images of T_ang_ cells and HUVEC. **(E)** Fold change of median loop area of individual donors from PWH and PWoH and HIVGag-specific CD8 T cells. **(F)** Fold change of median loop area of pooled all donors. HUVEC shows mean±SEM of pooled experiments with all donors. The bar graph is represented by box and whisker showing the median value with first and third quartiles in the box, with whiskers extending to the minimum and maximum values. Line graphs represent the average of all donors. Statistical analysis was performed using non-parametric Mann-Whitney test. *P* value < 0.05 was considered significant.

These observations suggest that secretion of cytokines and effector molecules from activated CD8 T_ang_ cells can lead to endothelial activation promoting trafficking of immune cells and secretion of proinflammatory and pro-angiogenic cytokines by endothelial cells.

We next performed a tube forming assay in which endothelial cells cultured in an extracellular matrix form capillary-like structures as a measurement of angiogenesis to better evaluate the interactions between CD8 T_ang_-endothelial cells (94). CD8 T_angs_ cells were sorted from PBMCs, labeled with CFSE and cultured with HUVEC at 1:2 T cell to HUVEC ratio (Figure 5C and D).

CD8 T_ang_ cells overtime migrate and adhere to the endothelial cells as the tubes formed showing a mesh of capillary-like structures and CFSE^+^CD8 T_ang_, suggesting a close interaction with endothelial cells in the extracellular matrix (Figure 5C and D). Overtime, CSFE^+^CD8 T_ang_ from PWH and sorted HIV_Gag_ specific CD8 T cells showed a trend of slowing the growth of the tube (loop area), although these responses were heterogenous among the donors (Figure 5E). In contrast, CD8 T_ang_ from PWoH showed a trend promoting bigger loop area overtime.

Altogether, these data imply a crosstalk between CD8 T_ang_ cells and endothelial cells. CD8 T_ang_ cells from PWH promote endothelial activation (ICAM1 and VCAM1), and induced secretion of proinflammatory and pro-angiogenic cytokines influencing endothelial cell function.

### Activated CD8 T_ang_ cells mediate endothelial cell activation through a granzyme-dependent PAR1 activation

We next investigated the mechanisms by which CD8 T_ang_ cells activate endothelial cells. PAR1 is expressed by endothelial cells and its signaling mediates important functions in maintaining barrier function (35). In addition to thrombin, PAR1 is activated by other proteases including extracellular granzymes (A, B and K) regulating angiogenesis and extracellular remodeling (51–53). These observations led us to the hypothesis that granzymes are involved in endothelial activation in the context of HIV infection.

We next studied the effects of granzymes in endothelial cells. To isolate the effects of granzymes from cytokines secreted by CD8 T_ang_ cells we used recombinant human (rh) GZMA and GZMK. GZMA and GZMK have tryptase activity and preferentially cleave after basic residues (Arg and Lys) which is similar to thrombin. GZMB is an aspartase and cleaves after Asp (48, 49, 54, 55).

We determined the effects of GZs dependent PAR1 activation on calcium mobilization. HUVEC were cultured *in vitro* and activated either with rhGZMA (100nM) and Thrombin (3nM). Compared to thrombin, GZMA induced a calcium mobilization with slower kinetics and sustained overtime (Figure 6B). These effects were not observed on GZMK and GZMB. We then investigated the downstream effects induced by GZM-dependent activation of PAR1 at later time points of stimulation and measure cytokine secretion by endothelial cells (Figure 6C).

**Figure 6.**
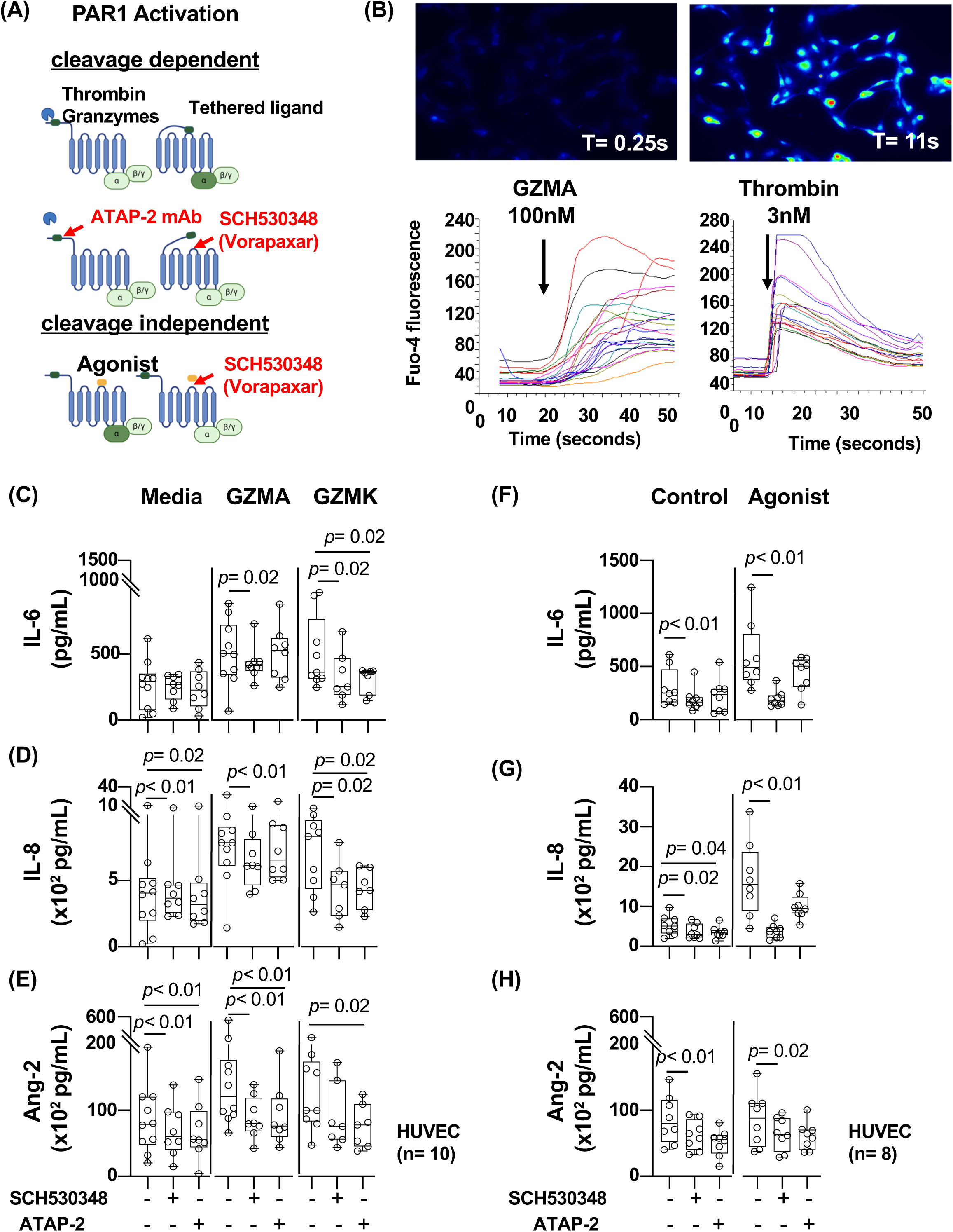
Activated CD8 T_ang_ cells mediate endothelial cell activation through a granzyme-dependent PAR1 activation. **(A)** Diagram illustrates the molecular mechanism of cleavage dependent PAR1 activation via serine proteases. Inhibition of PAR1 activation by ATAP-2 mAb and antagonist SCH530348 (vorapaxar). Peptide agonist (mimics the tethered ligand) and cleavage independent PAR1 activation. The figure was generated with Biorender. **(B)** Image of Granzyme A induced intracellular calcium mobilization in HUVEC. Intracellular calcium mobilization kinetics in HUVEC stimulated with granzyme A and thrombin as control. Each line represents the kinetics overtime of individual cells (lower panel). **(C, D and E)** HUVEC were stimulated with 100 nM of rh GZMA and GZMK overnight in the presence or absence of 10 μg/mL of ATAP-2 mAb or 300 nM of PAR1 antagonist SCH530348. (**F, G** and **H**) HUVEC were stimulated with 100 μM of PAR1 agonist peptide (TFLLR-NH2) and peptide control (RLLFT-NH2) as positive and negative controls respectively. IL-6, IL-8/CXCL8 and Angiopoietin-2 were measured in the supernatants of the cultures. The bar graph is represented by box and whisker showing the median value with first and third quartiles in the box, with whiskers extending to the minimum and maximum values. Statistical analysis was performed using non-parametric Mann-Whitney test. *P* value <0.05 was considered significant.

HUVEC were activated with either rhGZMA and rhGZMK (100 nM) in the absence or presence of PAR1 antagonist SCH530348 which inhibit binding of the tethered ligand (Figure 6A). In addition, we used the mAb ATAP-2 that blocks PAR1 cleavage by binding to an epitope within the PAR1 tethered ligand domain, Figure 6A (42, 95–97). As positive control, we used an agonist peptide specific for PAR1 (mimics the tethered ligand) and mediates activation of PAR1 in a cleavage independent manner (Figure 6A).

We found that rhGZMA and rhGZMK induced secretion of proinflammatory cytokines IL-6, IL-8/CXCL8 and Ang-2 (Figure 6 and Figure S6C). Moreover, GZA and GZK induced pro-angiogenic cytokine VEGF, albeit at lower levels (Figure S7). The antagonist SCH530348 (vorapaxar) partially blocked PAR1 activation by rhGZMA and consequently the secretion of IL-6 and IL-8 and Ang-2 (Figure 6C, D and E). The effects mediated by the ATAP-2 mAb showed inhibition Ang-2 but not IL-6 and IL-8.

In addition, rhGZMK dependent PAR1 activation by the antagonist SCH530348 inhibited secretion of IL-6 and IL-8/CXCL8 but not Ang-2 secretion by endothelial cells (Figure 6C, D and E). In contrast, ATAP-2 mAb blocked GZMK dependent PAR1 activation and the secretion of IL-6, IL-8/CXCL8 and Ang-2 (Figure 6C and S7). As expected, stimulation with PAR1 agonist peptide (cleavage independent activation) induced secretion of IL-6 and IL-8, however it did not induce a significant increase in secretion of Ang-2 (Figure 6H and S6C).

All together these data suggest that CD8 T_ang_ cells mediate endothelial cell activation in part through GZMA and GZMK dependent PAR1 activation leading to the secretion of proinflammatory cytokines.

## Discussion

Chronic CD8 T cell activation is associated with increased cardiovascular disease risk in PWH, however the mechanisms remain to be defined (6, 10, 16, 18, 22). Because endothelial inflammation and dysfunction are the main features of cardiovascular disease, in the present study we investigated a physiological mechanism in which the immune system (with a focus on CD8 T cells) intersects with vascular repair. PAR1 connects the coagulation cascade, endothelial cells and CD8 T cells at the site of vascular injury, in this study, using a PAR1 reporter mouse we established the kinetics of PAR1 expression in virus-specific CD8 T cells during the course of an acute viral infection with LCMV. PAR1 expression in LCMV-specific CD8 T cells is downregulated during the effector phase and its expression is upregulated during the contraction phase and early memory development (day 21 p.i). In addition, we identified a subset of CD31^high^CD8 T cells that secrete VEGF providing evidence that CD8 T_ang_ cells may contributes to vascular repair after viral control. T_ang_ cells were originally discovered in the center of endothelial progenitor cell colonies from PBMCs by Hur J. et al. (25–27). However, whether T_ang_ cells play a role during infection is not well understood. We found that expansion of CD31^high^CD8 T_ang_ cells occurs concomitant with the EC recovery and secrete proangiogenic cytokines VEGF and GMCSF. These cytokines are associated with endothelial cell proliferation, survival and bone marrow mobilization of endothelial progenitors (70–72, 98). Our study shows that CD31^high^CD8 T_ang_ cells may regulate angiogenesis by promoting the recruitment of endothelial progenitor cells, and other cells including myeloid cells to the site of vascular injury.

Particularly, CD31^high^CD8 T_ang_ cells expressed CD309 (VEGFR2) and secrete proangiogenic and Th1 cytokines. Similar to these findings, a study reported that hypoxia upregulates expression of CD309 (VEGFR2) and VEGF signaling, which in turn increased expression of IFNγ and inhibited IL-10 (71). In our model of LCMV infection, there is a mild endothelial injury that may induce hypoxia driving expression of CD309 and secretion of VEGF by CD8 T_ang_ cells. It is also possible VEGF signal promotes the acquisition of a proangiogenic property by CD8 T_ang_ cells and warrants further investigation.

CD8 T cells are predominantly viewed as main players in immunity against viruses and cancer due to their cytotoxic function, however, emerging evidence suggest that they play a role in tissue regeneration and remodeling. CD31^high^CD8 T_ang_ cells were LCMV-specific and secrete VEGF, IFNγ and TNFα and degranulate upon *in vitro* stimulation LCMVGp33 suggesting may acquire this function after viral control. More importantly, CD31^high^CD8 T_ang_ cells showed a memory precursor phenotype (KLGRG1^-^CD127^+^) suggesting a survival advantage during the contraction phase and a proliferative potential (66–69).

Expression of CD31 in T cells has been associated with T cell trafficking and studies have shown that CD31 attenuates TCR signaling in T cells as contains two immunoreceptor tyrosine-based inhibitory motif (ITIM) units in the cytoplasmic domain (99–101). It is possible that CD31 expression by CD8 T_ang_ cells allows close interactions with endothelial cells preventing their potential cytotoxic activity on endothelial cells. CD8 T cells with tissue remodeling properties that secrete IFNγ, TNFα and epidermal growth factor receptor (EGFR)-ligand amphiregulin (AREG) have been described highlighting CD8 T cell function beyond cytotoxicity (102). Moreover, in pathological conditions such as retinal vascular diseases, VEGF producing CD8 T cells drive inflammation and abnormal angiogenesis suggesting that dysregulated T_ang_ cell function may contribute to the pathology of disease (103).

We also investigated changes in lung endothelial cells. ECs are heterogenous and play distinct roles during immune response by supporting trafficking of immune cells, immune functions (immune cell-like ECs) and repair (regenerative ECs) (76, 104). Particularly, our study shows that at the peak of CD8 T cell recruitment into the lung, ECs expressed high levels of PD-L1 likely induced by cytokines produced by LCMV-specific CD8 T cells regulating the immune mediated-pathology including endothelial injury (105). The “immune-EC” phenotype returns to basal levels when the virus is controlled and ECs expressing receptors associated with cell proliferation and survival (CD34, CD309, CD105, CD201) are more predominant. Studies of lung endothelial cells in response to acute lung injury by LPS and influenza infection using single cell RNAseq, revealed distinct functions of ECs “immune-ECs” and “regenerative-ECs” the later involved in vascular remodeling suggesting an important cross-talk between ECs, host immunity and endothelial repair (77, 79). In contrast to these studies that used an acute respiratory lung viral infection or injury, we used a systemic virus to evaluate vascular injury and investigate potential mechanisms associated with chronic HIV infection and endothelial inflammation. Similar to our studies, the authors observed a trend of reduced EC count at day 7 p.i, (79). Although this study did not investigate the potential mechanisms driving proliferation of ECs in the reconstitution phase (between day 14 to 21 p.i), they described a proliferating population of CD34^+^ECs with regenerative capacity (79). We found that CD8 T_ang_ cells are associated with population of ECs (cluster 6) that express CD34 and have the potential to proliferate in response to proangiogenic cytokines through CD309 and CD105 signaling. CD309, the receptor for VEGF, is a critical factor for endothelial cell proliferation, migration, and survival, and plays important role during vascular remodeling (106). In addition, CD105, a marker for endothelial cell proliferation, is expressed on vascular endothelium during angiogenesis and prevents endothelial cell apoptosis induced by TGFβ (107). Moreover, this EC population express of endothelial protein C receptor (CD201) that has been proposed as a marker of endothelial progenitor cells with new vessel forming properties (86, 108).

T_ang_ cells are key players in maintaining endothelial homeostasis and alterations in vascular repair mechanisms may contribute to endothelial dysfunction and increase cardiovascular disease risk (109). In fact, reduced proportion of circulating T_angs_ has been associated with age and increased cardiovascular risk (25, 28, 29). Patients with rheumatoid arthritis and inflammatory diseases that suffered a cardiovascular event complication had lower circulating T_ang_ cells (29, 89, 110–114). In this study, we found that compared with uninfected controls (PWoH), CD8 T_ang_ cells from PWH had proinflammatory phenotype, express PAR1, and the adhesion and homing receptors CD49f and CX3CR1 respectively (32, 90, 115). In addition, CD8 T_ang_ cells had cytotoxic potential and expressed GZMA and GZMB. The activated phenotype of CD8 T_ang_ cells but not CD4 T_ang_ was associated with the ASCVD suggesting a relationship between endothelial inflammation and CD8 T_ang_ cell immune activation.

Granzymes (GZMA, GZMB, GZMK), can activate PAR1 and induce a “biased” or non-canonical signal distinct from thrombin (51, 116, 117). GZMs were first thought to be involved in cytotoxic function and recent evidence suggests non-cytotoxic and extracellular biological activities including regulation of vascular integrity, angiogenesis, extracellular matrix remodeling and trans-endothelial migration (45–47, 49, 51–53, 116, 118). The present study shows that, *in vitro*, CD8 T_ang_ cells induced activation of ECs via granzyme induced production of proinflammatory cytokines. In addition, granzyme dependent PAR1 activation provides a potential mechanism by which HIV alters proangiogenic function and the chronic stimulation of this pathway may lead endothelial inflammation and dysfunction.

Granzymes are emerging as potential contributors of inflammation in cardiovascular diseases (51, 116, 119, 120). Recent study reported a subset of CD8 T cells expression GZMK and GZMB with proinflammatory (rather than cytotoxic) function in chronically inflamed human tissues supporting their potential role in inflammation (53). Moreoever, increased levels of GZMs in biological fluids and tissues has been reported in inflammatory conditions including cardiovascular disease, autoimmunity, chronic obstructive pulmonary disease, and cancer (51, 53, 121, 122). Supporting these observations, we found an association between plasma GZMA leves and endothelial inflammation markers including sVCAM1 and IL-6 was observed in PWH suggesting that ongoing endothelial inflammation in the setting of suppressed viremia may be driven by T_ang_ cell activation.

Alltoghether these data highlight a novel role of CD8 T_ang_ cells in vascular remodeling, and granzyme-PAR1 dependent pathway promoting endothelial inglammation. CD8 T_ang_ cells and GZMA dependent PAR1 activation could be a target of intervetion to reduce endothelial inflammation and injury in PWH potentially mitigating the risk of cardiovascular disease during chronic HIV infection.

## Acknowledgements

Research reported in this publication was supported by the National Institute of Allergy and Infectious Diseases of the National Institutes of Health under award number NIH R01AI145549 and R21AI186960. We thank the District of Columbia Center for AIDS Research, an NIH funded program (P30AI117970) which is supported by the following NIH Co-Funding and Participating Institutes and Centers: NIAID, NCI, NICHD, NHLBI, NIDA, NIMH, NIA, NIDDK, NIMHD, NIDCR, NINR, FIC and OAR. The content of this publication does not necessarily reflect the views or policies of the Department of Health and Human Services, nor does mention of trade names, commercial products, or organizations imply endorsement by the U.S. Government.

## Authors’ contributions

MC conceptualized the study and supervise data analysis. TL, CM, CJ performed and analyzed mice experiments, JC, ZZ, MFC, MA, SY, performed human experiments, JK, PK, HC recruited patients, KDV, GPA advise and performed endothelial assay, DBG and LP advice mice infection model. MC, TL and LP wrote the manuscript.

## References

1. Baker JV, Sharma S, Grund B, Rupert A, Metcalf JA, Schechter M, et al. Systemic Inflammation, Coagulation, and Clinical Risk in the START Trial. Open Forum Infect Dis. 2017;4(4):ofx262.

2. Hadigan C, Paules CI, and Fauci AS. Association Between Human Immunodeficiency Virus Infection and Cardiovascular Diseases: Finding a Solution to Double Jeopardy. JAMA Cardiol. 2017;2(2):123–4.

3. Hsue PY, Deeks SG, and Hunt PW. Immunologic basis of cardiovascular disease in HIV-infected adults. The Journal of infectious diseases. 2012;205 Suppl 3:S375–82.

4. Sinha A, Ma Y, Scherzer R, Hur S, Li D, Ganz P, et al. Role of T-Cell Dysfunction, Inflammation, and Coagulation in Microvascular Disease in HIV. Journal of the American Heart Association. 2016;5(12).

5. Freiberg MS, Chang CC, Kuller LH, Skanderson M, Lowy E, Kraemer KL, et al. HIV infection and the risk of acute myocardial infarction. JAMA internal medicine. 2013;173(8):614–22.

6. Mattingly AS, Unsal AB, Purdy JB, Gharib AM, Rupert A, Kovacs JA, et al. T-cell Activation and E-selectin Are Associated With Coronary Plaque in HIV-infected Young Adults. The Pediatric infectious disease journal. 2017;36(1):63–5.

7. Serrano-Villar S, Sainz T, Lee SA, Hunt PW, Sinclair E, Shacklett BL, et al. HIV-infected individuals with low CD4/CD8 ratio despite effective antiretroviral therapy exhibit altered T cell subsets, heightened CD8+ T cell activation, and increased risk of non-AIDS morbidity and mortality. PLoS Pathog. 2014;10(5):e1004078.

8. Stein JH, and Hsue PY. Inflammation, immune activation, and CVD risk in individuals with HIV infection. JAMA. 2012;308(4):405–6.

9. Triant VA, Perez J, Regan S, Massaro JM, Meigs JB, Grinspoon SK, et al. Cardiovascular Risk Prediction Functions Underestimate Risk in HIV Infection. Circulation. 2018;137(21):2203–14.

10. Shah ASV, Stelzle D, Lee KK, Beck EJ, Alam S, Clifford S, et al. Global Burden of Atherosclerotic Cardiovascular Disease in People Living With HIV: Systematic Review and Meta-Analysis. Circulation. 2018;138(11):1100–12.

11. Lundgren JD, Babiker AG, Sharma S, Grund B, Phillips AN, Matthews G, et al. Long-Term Benefits from Early Antiretroviral Therapy Initiation in HIV Infection. NEJM Evid. 2023;2(3).

12. Catalfamo M, Wilhelm C, Tcheung L, Proschan M, Friesen T, Park JH, et al. CD4 and CD8 T cell immune activation during chronic HIV infection: roles of homeostasis, HIV, type I IFN, and IL-7. J Immunol. 2011;186(4):2106–16.

13. Nou E, Lo J, and Grinspoon SK. Inflammation, immune activation, and cardiovascular disease in HIV. *AIDS (London*, England). 2016;30(10):1495–509.

14. Imamichi H, Smith M, Adelsberger JW, Izumi T, Scrimieri F, Sherman BT, et al. Defective HIV-1 proviruses produce viral proteins. Proceedings of the National Academy of Sciences of the United States of America. 2020;117(7):3704–10.

15. Benlarbi M, Richard J, Bourassa C, Tolbert WD, Chartrand-Lefebvre C, Gendron-Lepage G, et al. Plasmatic HIV-1 soluble gp120 is associated with correlates of immune dysfunction and inflammation in ART-treated individuals with undetectable viremia. The Journal of infectious diseases. 2023.

16. Longenecker CT, Funderburg NT, Jiang Y, Debanne S, Storer N, Labbato DE, et al. Markers of inflammation and CD8 T-cell activation, but not monocyte activation, are associated with subclinical carotid artery disease in HIV-infected individuals. HIV Med. 2013;14(6):385–90.

17. Hsue PY, Deeks SG, and Hunt PW. Immunologic basis of cardiovascular disease in HIV-infected adults. The Journal of infectious diseases. 2012;205 Suppl 3(Suppl 3):S375–82.

18. Kaplan RC, Sinclair E, Landay AL, Lurain N, Sharrett AR, Gange SJ, et al. T cell activation and senescence predict subclinical carotid artery disease in HIV-infected women. J Infect Dis. 2011;203(4):452–63.

19. McLaughlin MM, Ma Y, Scherzer R, Rahalkar S, Martin JN, Mills C, et al. Association of Viral Persistence and Atherosclerosis in Adults With Treated HIV Infection. JAMA Netw Open. 2020;3(10):e2018099.

20. Turcotte I, El-Far M, Sadouni M, Chartrand-Lefebvre C, Filali-Mouhim A, Fromentin R, et al. Association Between the Development of Subclinical Cardiovascular Disease and Human Immunodeficiency Virus (HIV) Reservoir Markers in People With HIV on Suppressive Antiretroviral Therapy. Clin Infect Dis. 2023;76(7):1318–21.

21. Wang Y, Brichacek B, Dubrovsky L, Pushkarsky T, Korolowicz K, Rodriguez O, et al. Increased Atherosclerosis in HIV-Infected Humanized Mice Is Caused by a Single Viral Protein, Nef. The Journal of infectious diseases. 2025;232(1):e116–e25.

22. Karim R, Mack WJ, Kono N, Tien PC, Anastos K, Lazar J, et al. T-cell activation, both pre- and post-HAART levels, correlates with carotid artery stiffness over 6.5 years among HIV-infected women in the WIHS. J Acquir Immune Defic Syndr. 2014;67(3):349–56.

23. Keller TT, Mairuhu AT, de Kruif MD, Klein SK, Gerdes VE, ten Cate H, et al. Infections and endothelial cells. Cardiovascular research. 2003;60(1):40–8.

24. Wang Y, Brichacek B, Dubrovsky L, Pushkarsky T, Korolowicz K, Rodriguez O, et al. Increased atherosclerosis in HIV-infected humanized mice is caused by a single viral protein, Nef. The Journal of infectious diseases. 2025.

25. Hur J, Yang HM, Yoon CH, Lee CS, Park KW, Kim JH, et al. Identification of a novel role of T cells in postnatal vasculogenesis: characterization of endothelial progenitor cell colonies. Circulation. 2007;116(15):1671–82.

26. Asahara T, Murohara T, Sullivan A, Silver M, van der Zee R, Li T, et al. Isolation of putative progenitor endothelial cells for angiogenesis. *Science (New York*, NY. 1997;275(5302):964-7.

27. Yoder MC. Endothelial stem and progenitor cells (stem cells): (2017 Grover Conference Series). Pulm Circ. 2018;8(1):2045893217743950.

28. Rouhl RP, Mertens AE, van Oostenbrugge RJ, Damoiseaux JG, Debrus-Palmans LL, Henskens LH, et al. Angiogenic T-cells and putative endothelial progenitor cells in hypertension-related cerebral small vessel disease. Stroke; a journal of cerebral circulation. 2012;43(1):256–8.

29. Rodriguez-Carrio J, Alperi-Lopez M, Lopez P, Alonso-Castro S, Ballina-Garcia FJ, and Suarez A. Angiogenic T cells are decreased in rheumatoid arthritis patients. Annals of the rheumatic diseases. 2015;74(5):921–7.

30. Hurley A, Smith M, Karpova T, Hasley RB, Belkina N, Shaw S, et al. Enhanced effector function of CD8(+) T cells from healthy controls and HIV-infected patients occurs through thrombin activation of protease-activated receptor 1. The Journal of infectious diseases. 2013;207(4):638–50.

31. Green SA, Smith M, Hasley RB, Stephany D, Harned A, Nagashima K, et al. Activated platelet-T-cell conjugates in peripheral blood of patients with HIV infection: coupling coagulation/inflammation and T cells. *AIDS (London*, England). 2015;29(11):1297–308.

32. Mudd JC, Panigrahi S, Kyi B, Moon SH, Manion MM, Younes SA, et al. Inflammatory Function of CX3CR1+ CD8+ T Cells in Treated HIV Infection Is Modulated by Platelet Interactions. The Journal of infectious diseases. 2016;214(12):1808–16.

33. Chen H, Smith M, Herz J, Li T, Hasley R, Le Saout C, et al. The role of protease-activated receptor 1 signaling in CD8 T cell effector functions. iScience. 2021;24(11):103387.

34. Coughlin SR. How the protease thrombin talks to cells. Proceedings of the National Academy of Sciences of the United States of America. 1999;96(20):11023–7.

35. Coughlin SR. Thrombin signalling and protease-activated receptors. Nature. 2000;407(6801):258–64.

36. Soh UJ, Dores MR, Chen B, and Trejo J. Signal transduction by protease-activated receptors. British journal of pharmacology. 2010;160(2):191–203.

37. Sambrano GR, Weiss EJ, Zheng YW, Huang W, and Coughlin SR. Role of thrombin signalling in platelets in haemostasis and thrombosis. Nature. 2001;413(6851):74-8.

38. Kahn ML, Zheng YW, Huang W, Bigornia V, Zeng D, Moff S, et al. A dual thrombin receptor system for platelet activation. Nature. 1998;394(6694):690-4.

39. Ishihara H, Zeng D, Connolly AJ, Tam C, and Coughlin SR. Antibodies to protease-activated receptor 3 inhibit activation of mouse platelets by thrombin. Blood. 1998;91(11):4152–7.

40. Xu WF, Andersen H, Whitmore TE, Presnell SR, Yee DP, Ching A, et al. Cloning and characterization of human protease-activated receptor 4. Proceedings of the National Academy of Sciences of the United States of America. 1998;95(12):6642–6.

41. Hollenberg MD. PARs in the stars: proteinase-activated receptors and astrocyte function. Focus on “Thrombin (PAR-1)-induced proliferation in astrocytes via MAPK involves multiple signaling pathways”. American journal of physiology Cell physiology. 2002;283(5):C1347–50.

42. Ramachandran R, Noorbakhsh F, Defea K, and Hollenberg MD. Targeting proteinase-activated receptors: therapeutic potential and challenges. Nature reviews Drug discovery. 2012;11(1):69–86.

43. Wang T, Lee MH, Choi E, Pardo-Villamizar CA, Lee SB, Yang IH, et al. Granzyme B-induced neurotoxicity is mediated via activation of PAR-1 receptor and Kv1.3 channel. PLoS One. 2012;7(8):e43950.

44. Zhao P, Metcalf M, and Bunnett NW. Biased signaling of protease-activated receptors. Frontiers in endocrinology. 2014;5:67.

45. Lee PR, Johnson TP, Gnanapavan S, Giovannoni G, Wang T, Steiner JP, et al. Protease-activated receptor-1 activation by granzyme B causes neurotoxicity that is augmented by interleukin-1beta. J Neuroinflammation. 2017;14(1):131.

46. Cooper DM, Pechkovsky DV, Hackett TL, Knight DA, and Granville DJ. Granzyme K activates protease-activated receptor-1. PLoS One. 2011;6(6):e21484.

47. Sharma M, Merkulova Y, Raithatha S, Parkinson LG, Shen Y, Cooper D, et al. Extracellular granzyme K mediates endothelial activation through the cleavage of protease-activated receptor-1. The FEBS journal. 2016;283(9):1734–47.

48. Bovenschen N, Quadir R, van den Berg AL, Brenkman AB, Vandenberghe I, Devreese B, et al. Granzyme K displays highly restricted substrate specificity that only partially overlaps with granzyme A. J Biol Chem. 2009;284(6):3504–12.

49. Suidan HS, Bouvier J, Schaerer E, Stone SR, Monard D, and Tschopp J. Granzyme A released upon stimulation of cytotoxic T lymphocytes activates the thrombin receptor on neuronal cells and astrocytes. Proc Natl Acad Sci U S A. 1994;91(17):8112–6.

50. Kaiserman D, Zhao P, Rowe CL, Leong A, Barlow N, Joeckel LT, et al. Granzyme K initiates IL-6 and IL-8 release from epithelial cells by activating protease-activated receptor 2. PLoS One. 2022;17(7):e0270584.

51. Zeglinski MR, and Granville DJ. Granzymes in cardiovascular injury and disease. Cellular signalling. 2020;76:109804.

52. Arias M, Martinez-Lostao L, Santiago L, Ferrandez A, Granville DJ, and Pardo J. The Untold Story of Granzymes in Oncoimmunology: Novel Opportunities with Old Acquaintances. Trends Cancer. 2017;3(6):407–22.

53. Jonsson AH, Zhang F, Dunlap G, Gomez-Rivas E, Watts GFM, Faust HJ, et al. Granzyme K(+) CD8 T cells form a core population in inflamed human tissue. Sci Transl Med. 2022;14(649):eabo0686.

54. Mahrus S, and Craik CS. Selective chemical functional probes of granzymes A and B reveal granzyme B is a major effector of natural killer cell-mediated lysis of target cells. Chem Biol. 2005;12(5):567–77.

55. Plasman K, Demol H, Bird PI, Gevaert K, and Van Damme P. Substrate specificities of the granzyme tryptases A and K. J Proteome Res. 2014;13(12):6067–77.

56. !!! INVALID CITATION !!! (52).

57. Harker JA, Wong KA, Dallari S, Bao P, Dolgoter A, Jo Y, et al. Interleukin-27R Signaling Mediates Early Viral Containment and Impacts Innate and Adaptive Immunity after Chronic Lymphocytic Choriomeningitis Virus Infection. Journal of virology. 2018;92(12).

58. Haghverdi L, Lun ATL, Morgan MD, and Marioni JC. Batch effects in single-cell RNA-sequencing data are corrected by matching mutual nearest neighbors. Nat Biotechnol. 2018;36(5):421–7.

59. Anderson KG, Sung H, Skon CN, Lefrancois L, Deisinger A, Vezys V, et al. Cutting edge: intravascular staining redefines lung CD8 T cell responses. J Immunol. 2012;189(6):2702–6.

60. Masopust D, Vezys V, Marzo AL, and Lefrancois L. Preferential localization of effector memory cells in nonlymphoid tissue. Science (New York, NY. 2001 ;291(5512):2413-7.

61. Purton JF, Martin CE, and Surh CD. Enhancing T cell memory: IL-7 as an adjuvant to boost memory T-cell generation. Immunol Cell Biol. 2008;86(5):385–6.

62. Kang SS, and McGavern DB. Lymphocytic choriomeningitis infection of the central nervous system. Frontiers in bioscience : a journal and virtual library. 2008;13:4529–43.

63. Baccala R, Welch MJ, Gonzalez-Quintial R, Walsh KB, Teijaro JR, Nguyen A, et al. Type I interferon is a therapeutic target for virus-induced lethal vascular damage. Proceedings of the National Academy of Sciences of the United States of America. 2014;111(24):8925–30.

64. Wherry EJ, Teichgraber V, Becker TC, Masopust D, Kaech SM, Antia R, et al. Lineage relationship and protective immunity of memory CD8 T cell subsets. Nature immunology. 2003;4(3):225–34.

65. Wherry EJ, Blattman JN, Murali-Krishna K, van der Most R, and Ahmed R. Viral persistence alters CD8 T-cell immunodominance and tissue distribution and results in distinct stages of functional impairment. Journal of virology. 2003;77(8):4911–27.

66. Joshi NS, Cui W, Chandele A, Lee HK, Urso DR, Hagman J, et al. Inflammation directs memory precursor and short-lived effector CD8(+) T cell fates via the graded expression of T-bet transcription factor. Immunity. 2007;27(2):281–95.

67. Sarkar S, Kalia V, Haining WN, Konieczny BT, Subramaniam S, and Ahmed R. Functional and genomic profiling of effector CD8 T cell subsets with distinct memory fates. The Journal of experimental medicine. 2008;205(3):625–40.

68. Chang JT, Wherry EJ, and Goldrath AW. Molecular regulation of effector and memory T cell differentiation. Nature immunology. 2014;15(12):1104–15.

69. Herndler-Brandstetter D, Ishigame H, Shinnakasu R, Plajer V, Stecher C, Zhao J, et al. KLRG1(+) Effector CD8(+) T Cells Lose KLRG1, Differentiate into All Memory T Cell Lineages, and Convey Enhanced Protective Immunity. Immunity. 2018;48(4):716–29 e8.

70. Perez-Gutierrez L, and Ferrara N. Biology and therapeutic targeting of vascular endothelial growth factor A. Nat Rev Mol Cell Biol. 2023;24(11):816–34.

71. Mor F, Quintana FJ, and Cohen IR. Angiogenesis-inflammation cross-talk: vascular endothelial growth factor is secreted by activated T cells and induces Th1 polarization. J Immunol. 2004;172(7):4618–23.

72. Wang QR, Wang F, Zhu WB, Lei J, Huang YH, Wang BH, et al. GM-CSF accelerates proliferation of endothelial progenitor cells from murine bone marrow mononuclear cells in vitro. Cytokine. 2009;45(3):174–8.

73. Bussolino F, Wang JM, Defilippi P, Turrini F, Sanavio F, Edgell CJ, et al. Granulocyte- and granulocyte-macrophage-colony stimulating factors induce human endothelial cells to migrate and proliferate. Nature. 1989;337(6206):471-3.

74. Biedermann BC. Vascular endothelium: checkpoint for inflammation and immunity. News in physiological sciences : an international journal of physiology produced jointly by the International Union of Physiological Sciences and the American Physiological Society. 2001;16:84–8.

75. Danese S, Dejana E, and Fiocchi C. Immune regulation by microvascular endothelial cells: directing innate and adaptive immunity, coagulation, and inflammation. J Immunol. 2007;178(10):6017–22.

76. Amersfoort J, Eelen G, and Carmeliet P. Immunomodulation by endothelial cells - partnering up with the immune system? Nature reviews. 2022;22(9):576–88.

77. Zhang L, Gao S, White Z, Dai Y, Malik AB, and Rehman J. Single-cell transcriptomic profiling of lung endothelial cells identifies dynamic inflammatory and regenerative subpopulations. JCI Insight. 2022;7(11).

78. Godoy RS, Cober ND, Cook DP, McCourt E, Deng Y, Wang L, et al. Single-cell transcriptomic atlas of lung microvascular regeneration after targeted endothelial cell ablation. Elife. 2023;12.

79. Niethamer TK, Stabler CT, Leach JP, Zepp JA, Morley MP, Babu A, et al. Defining the role of pulmonary endothelial cell heterogeneity in the response to acute lung injury. Elife. 2020;9.

80. Rodor J, Chen SH, Scanlon JP, Monteiro JP, Caudrillier A, Sweta S, et al. Single-cell RNA sequencing profiling of mouse endothelial cells in response to pulmonary arterial hypertension. Cardiovasc Res. 2022;118(11):2519–34.

81. Jerkic M, Rivas-Elena JV, Prieto M, Carron R, Sanz-Rodriguez F, Perez-Barriocanal F, et al. Endoglin regulates nitric oxide-dependent vasodilatation. FASEB journal : official publication of the Federation of American Societies for Experimental Biology. 2004;18(3):609–11.

82. Toporsian M, Gros R, Kabir MG, Vera S, Govindaraju K, Eidelman DH, et al. A role for endoglin in coupling eNOS activity and regulating vascular tone revealed in hereditary hemorrhagic telangiectasia. Circulation research. 2005;96(6):684–92.

83. Kretschmer S, Dethlefsen I, Hagner-Benes S, Marsh LM, Garn H, and Konig P. Visualization of intrapulmonary lymph vessels in healthy and inflamed murine lung using CD90/Thy-1 as a marker. PLoS One. 2013;8(2):e55201.

84. Levine JH, Simonds EF, Bendall SC, Davis KL, Amir el AD, Tadmor MD, et al. Data-Driven Phenotypic Dissection of AML Reveals Progenitor-like Cells that Correlate with Prognosis. Cell. 2015;162(1):184–97.

85. Li Q, Wei S, Li Y, Wu F, Qin X, Li Z, et al. Blocking of programmed cell death-ligand 1 (PD-L1) expressed on endothelial cells promoted the recruitment of CD8(+)IFN-gamma(+) T cells in atherosclerosis. Inflammation research : official journal of the European Histamine Research Society [et al]. 2023;72(4):783–96.

86. Yu QC, Song W, Wang D, and Zeng YA. Identification of blood vascular endothelial stem cells by the expression of protein C receptor. Cell Res. 2016;26(10):1079–98.

87. Thiyagarajan M, Cheng T, and Zlokovic BV. Endothelial cell protein C receptor: role beyond endothelium? Circulation research. 2007;100(2):155–7.

88. Zhu Z, Li T, Chen J, Kumar J, Kumar P, Qin J, et al. The Role of Inflammation and Immune Activation on Circulating Endothelial Progenitor Cells in Chronic HIV Infection. Frontiers in immunology. 2021;12:663412.

89. Kushner EJ, MacEneaney OJ, Morgan RG, Van Engelenburg AM, Van Guilder GP, and DeSouza CA. CD31+ T cells represent a functionally distinct vascular T cell phenotype. Blood Cells Mol Dis. 2010;44(2):74–8.

90. Pilling JE, Galvin A, Robins AM, Sewell HF, and Mahida YR. Expression of alpha5 (CD49e) and alpha6 (CD49f) integrin subunits on T cells in the circulation and the lamina propria of normal and inflammatory bowel disease colonic mucosa. Scandinavian journal of immunology. 1998;48(4):425–8.

91. Singh V, Kaur R, Kumari P, Pasricha C, and Singh R. ICAM-1 and VCAM-1: Gatekeepers in various inflammatory and cardiovascular disorders. Clin Chim Acta. 2023;548:117487.

92. Groten SA, Smit ER, Janssen EFJ, van den Eshof BL, van Alphen FPJ, van der Zwaan C, et al. Multi-omics delineation of cytokine-induced endothelial inflammatory states. Commun Biol. 2023;6(1):525.

93. Nakamichi M, Akishima-Fukasawa Y, Fujisawa C, Mikami T, Onishi K, and Akasaka Y. Basic Fibroblast Growth Factor Induces Angiogenic Properties of Fibrocytes to Stimulate Vascular Formation during Wound Healing. The American journal of pathology. 2016;186(12):3203–16.

94. Arnaoutova I, and Kleinman HK. In vitro angiogenesis: endothelial cell tube formation on gelled basement membrane extract. Nature protocols. 2010;5(4):628–35.

95. Zhang C, Srinivasan Y, Arlow DH, Fung JJ, Palmer D, Zheng Y, et al. High-resolution crystal structure of human protease-activated receptor 1. Nature. 2012;492(7429):387–92.

96. Brass LF, Pizarro S, Ahuja M, Belmonte E, Blanchard N, Stadel JM, et al. Changes in the structure and function of the human thrombin receptor during receptor activation, internalization, and recycling. J Biol Chem. 1994;269(4):2943–52.

97. O’Brien PJ, Prevost N, Molino M, Hollinger MK, Woolkalis MJ, Woulfe DS, et al. Thrombin responses in human endothelial cells. Contributions from receptors other than PAR1 include the transactivation of PAR2 by thrombin-cleaved PAR1. J Biol Chem. 2000;275(18):13502–9.

98. Qiu C, Xie Q, Zhang D, Chen Q, Hu J, and Xu L. GM-CSF induces cyclin D1 expression and proliferation of endothelial progenitor cells via PI3K and MAPK signaling. Cell Physiol Biochem. 2014;33(3):784–95.

99. Newman PJ. Switched at birth: a new family for PECAM-1. The Journal of clinical investigation. 1999;103(1):5–9.

100. Fornasa G, Groyer E, Clement M, Dimitrov J, Compain C, Gaston AT, et al. TCR stimulation drives cleavage and shedding of the ITIM receptor CD31. J Immunol. 2010;184(10):5485–92.

101. Ma L, Mauro C, Cornish GH, Chai JG, Coe D, Fu H, et al. Ig gene-like molecule CD31 plays a nonredundant role in the regulation of T-cell immunity and tolerance. Proceedings of the National Academy of Sciences of the United States of America. 2010;107(45):19461–6.

102. Delacher M, Schmidleithner L, Simon M, Stuve P, Sanderink L, Hotz-Wagenblatt A, et al. The effector program of human CD8 T cells supports tissue remodeling. The Journal of experimental medicine. 2024;221(2).

103. Deliyanti D, Figgett WA, Gebhardt T, Trapani JA, Mackay F, and Wilkinson-Berka JL. CD8(+) T Cells Promote Pathological Angiogenesis in Ocular Neovascular Disease. Arterioscler Thromb Vasc Biol. 2023;43(4):522–36.

104. Pober JS, and Sessa WC. Evolving functions of endothelial cells in inflammation. Nature reviews. 2007;7(10):803–15.

105. Frebel H, Nindl V, Schuepbach RA, Braunschweiler T, Richter K, Vogel J, et al. Programmed death 1 protects from fatal circulatory failure during systemic virus infection of mice. The Journal of experimental medicine. 2012;209(13):2485–99.

106. Lee C, Kim MJ, Kumar A, Lee HW, Yang Y, and Kim Y. Vascular endothelial growth factor signaling in health and disease: from molecular mechanisms to therapeutic perspectives. Signal Transduct Target Ther. 2025;10(1):170.

107. Li C, Issa R, Kumar P, Hampson IN, Lopez-Novoa JM, Bernabeu C, et al. CD105 prevents apoptosis in hypoxic endothelial cells. Journal of cell science. 2003;116(Pt 13):2677–85.

108. Lin Y, Banno K, Gil CH, Myslinski J, Hato T, Shelley WC, et al. Origin, prospective identification, and function of circulating endothelial colony-forming cells in mice and humans. JCI Insight. 2023;8(5).

109. Weil BR, Kushner EJ, Diehl KJ, Greiner JJ, Stauffer BL, and Desouza CA. CD31+ T cells, endothelial function and cardiovascular risk. Heart Lung Circ. 2011;20(10):659–62.

110. Lopez P, Rodriguez-Carrio J, Martinez-Zapico A, Caminal-Montero L, and Suarez A. Senescent profile of angiogenic T cells from systemic lupus erythematosus patients. J Leukoc Biol. 2016;99(3):405–12.

111. de Boer SA, Reijrink M, Abdulahad WH, Hoekstra ES, Slart R, Heerspink HJL, et al. Angiogenic T cells are decreased in people with type 2 diabetes mellitus and recruited by the dipeptidyl peptidase-4 inhibitor Linagliptin: A subanalysis from a randomized, placebo-controlled trial (RELEASE study). Diabetes Obes Metab. 2020;22(7):1220–5.

112. Kul A, Ozturk N, Kurt AK, and Arslan Y. Detection of Angiogenic T Cells and Endothelial Progenitor Cells in Behcet Disease and Determination of Their Relationship with Disease Activity. Life (Basel). 2023;13(6).

113. Manetti M, Pratesi S, Romano E, Bellando-Randone S, Rosa I, Guiducci S, et al. Angiogenic T cell expansion correlates with severity of peripheral vascular damage in systemic sclerosis. PLoS One. 2017;12(8):e0183102.

114. Zhang G, Liu Y, Qiu Y, Zhang J, Sun J, Zhou Z, et al. Circulating senescent angiogenic T cells are linked with endothelial dysfunction and systemic inflammation in hypertension. J Hypertens. 2021;39(5):970–8.

115. Panigrahi S, Chen B, Fang M, Potashnikova D, Komissarov AA, Lebedeva A, et al. CX3CL1 and IL-15 Promote CD8 T cell chemoattraction in HIV and in atherosclerosis. PLoS Pathog. 2020;16(9):e1008885.

116. Chamberlain CM, and Granville DJ. The role of Granzyme B in atheromatous diseases. Canadian journal of physiology and pharmacology. 2007;85(1):89–95.

117. Choy JC, McDonald PC, Suarez AC, Hung VH, Wilson JE, McManus BM, et al. Granzyme B in atherosclerosis and transplant vascular disease: association with cell death and atherosclerotic disease severity. Modern pathology : an official journal of the United States and Canadian Academy of Pathology, Inc. 2003;16(5):460–70.

118. Richardson KC, Jung K, Pardo J, Turner CT, and Granville DJ. Noncytotoxic Roles of Granzymes in Health and Disease. Physiology (Bethesda). 2022;37(6):323–48.

119. Ikemoto T, Hojo Y, Kondo H, Takahashi N, Hirose M, Nishimura Y, et al. Plasma granzyme B as a predicting factor of coronary artery disease--clinical significance in patients with chronic renal failure. Journal of cardiology. 2009;54(3):409–15.

120. Hendel A, Hiebert PR, Boivin WA, Williams SJ, and Granville DJ. Granzymes in age-related cardiovascular and pulmonary diseases. Cell death and differentiation. 2010;17(4):596–606.

121. Xu T, Zhu HX, You X, Ma JF, Li X, Luo PY, et al. Single-cell profiling reveals pathogenic role and differentiation trajectory of granzyme K+CD8+ T cells in primary Sjogren’s syndrome. JCI Insight. 2023;8(8).

122. Hiebert PR, Boivin WA, Abraham T, Pazooki S, Zhao H, and Granville DJ. Granzyme B contributes to extracellular matrix remodeling and skin aging in apolipoprotein E knockout mice. Experimental gerontology. 2011;46(6):489–99.

