## Supplemental Materials and Figures for "Angiogenic CD8 T cells from PWH induce Granzymes-dependent PAR1 activation promoting endothelial inflammation"

**Figure S1. Phenotypic characterization of lymphoid organs from naïve C57BL/6-F2r$^{fl/fl}$ mice.** **(A)** Scheme of the Exon 2 of F2r gene conditionally targeted. **(B)** Live cells and singlets were pre-gated followed by analysis of total hematopoietic cells (CD45$^{+}$) to assess cell counts in lymph nodes (LNs) and spleen from C57BL/6-F2r$^{fl/fl}$ (n= 5, open red symbol) and C57BL/6 (n= 5, open black symbol). **(C)** Gating strategy of immune cell subsets (CD45$^{+}$): NK cells (CD3$^{-}$NK1.1$^{+}$), T cells (CD3$^{+}$), Eosinophils (Eos, SSC-A$^{high}$Gr1$^{int}$), Neutrophils (Neut, Gr1$^{+}$), MHC-II$^{+}$ monocyte/macrophage (R1: B220$^{-}$MHC-II$^{+}$CD11b$^{+}$), B cells (R2: MHCII$^{+}$B220$^{+}$), plasmacytoid dendritic cells (pDCs) (R3: B220$^{+}$MHCII$^{-}$CD11c$^{+}$). MHCII$^{-}$ monocyte/macrophage (R4: B220$^{-}$MHCII$^{-}$CD11b$^{+}$), conventional dendritic cells (cDCs) (R1: B220$^{-}$MHCII$^{+}$CD11c$^{+}$). **(D)** Percentage and cell count of myeloid subsets (Eos, Neut, cDCs, pDCs and monocyte/macrophage) in LNs and spleen. **(E)** Percentage and cell count of lymphocytes B cells, NK cells and T cells in LNs and spleen. The bar graph is represented by box and whisker showing the median value with first and third quartiles in the box, with whiskers extending to the minimum and maximum values. Statistical analysis was performed using non-parametric Mann-Whitney test. *P* value <0.05 was considered significant.

**Figure S2. Expression of PAR1 in CD4 and CD8 T cells.**

**(**A) Splenic cells from C57BL/6-F2r$^{fl/fl}$ mice were isolated and activated *in vitro* with anti-CD3/CD28 mAbs for 3 days and expanded with IL-2 for an additional 5 days. PAR1 expression was assessed in resting and activated T cells by flow cytometry using rabbit anti-PAR1 mAb (Abcam, catal. ab322457) followed by an Alexa 647 labeled polyclonal anti-rabbit IgG. Contour plots of tdTomato and PAR1 expression by resting and *in vitro* activated CD4 and CD8 T cells.

**(**Lower panel**)** Histogram of PAR1 expression by resting and *in vitro* activated CD4 and CD8 T cells.

**Figure S3. PAR1 expression in lung ECs and Granzyme A expression by CD8 T cells in acute LCMV infection**

**(A)** C57BL/6-F2r$^{fl/fl}$ mice were infected with LCMV Armstrong (Arm) strain (2 x $10^5$ pfu, i.v). Day 6 and 21 p.i., splenic and lung immune cells were isolated and stimulated *in vitro* with LCMVGp33-41 peptide for 4 hours. DMSO was used as control. Representative contour plots of IFNγ and TNFα expression by CD8 T cells. Frequency of splenic and lung CD8 T cells secreting IFNγ and TNFα from naïve (N) mice and infected mice day 6 and day 21 p.i. **(B)** Representative contour plots of IFNγ and CD107a expression by CD8 T cells. Frequency of total (blue gate) CD107a$^+$ CD8 T cells in infected mice day 6 and day 21 p.i Naïve (N) animals were used as controls. **(C)** Median Fluorescence Intensity (MFI) of tdTomato expression in splenic and lung total CD8 T cells from naïve (N) and infected mice at day 6 and 21 p.i. **(D)** Representative contour plots of GMZA in degranulating (CD107a$^+$) LCMV-specific CD8 T cells from spleen and lung. at day 6 p.i. Naïve (N) animals were used as controls. **(E)** Median fluorescence intensity of PAR1 (tdT$^+$) expression in CD107a$^+$ and CD107$^-$ LCMV-specific CD8 T cells. The data in line graphs represent mean±SEM in each time point. Data are pooled from three independent experiments with 3-5 naïve animals and 5-17 per time point (F and H). The bar graph is represented by box and whisker showing the median value with first and third quartiles in the box, with whiskers extending to the minimum and maximum values. Statistical analysis was performed using non-parametric Mann-Whitney test. *P* value $< 0.05$ was considered significant.

**Figure S4. Expression of PAR1 in CD31$^{high}$CD8 T cells during LCMV infection.**

**(A)** C57BL/6-F2r$^{fl/fl}$ mice were infected with LCMV Armstrong (Arm) strain (2 x $10^5$ pfu, i.v). At day 15 p.i., mice were administered anti-CD8α mAb and IgG2b isotype for control mice. LCMV *glycoprotein* (*Gp*) mRNA expression was assessed in spleen and lung at day 6 and 21 p.i, and in mice treated with IgG and anti-CD8 mAb at day 21 p.i. Spleen collected from mice at day 3 p.i. was used as positive control (red dash line). **(B)** Representative contour plot of CD31$^{high}$ and CD309 expression by lung CD8 T cells from naïve mice and infected mice. Frequency and cell count of CD31$^{high}$CD8 T cells from naïve (N) mice and infected mice at day 6 and 21 p.i. Median fluorescence intensity of PAR1 (tdT$^{+}$) expression in CD31$^{high}$ and CD31$^{high}$ CD8 T cells. **(C)** Lung CD31$^{high}$CD8 and CD31$^{neg}$CD8 T cells LCMVArm infected C57BL/6-F2r$^{fl/fl}$ mice were sorted at day 21 p.i. CD31$^{high}$CD8 T cells were sorted from Naïve (N) mice (n= 3). Sorted T cells were stimulated *in vitro* in anti-CD3 and anti-CD28 mAbs coated plate. After 3 days stimulation cytokines were measured in the supernatants. Detection of growth factors macrophage colony-stimulating factor (M-CSF), platelet derived growth factor-A (PDGF-A) and granulocyte macrophage colony stimulating factor (GM-CSF), IL-2, IL-10, Th2 cytokines (IL-4, IL-5 and IL-13), Th17 cytokines (IL-17A and IL-6), Th22 cytokines (IL-22), Th9 (IL-9). The graph is represented by box and whisker showing the median value with first and third quartiles in the box, with whiskers extending to the minimum and maximum values. The data in line graphs represent mean±SEM in each time point. Data are pooled from two independent experiments with 3 naïve animals and 5-10 per time point. Statistical analysis was performed using non-parametric Mann-Whitney test. $P$ value < 0.05 was considered significant.

**Figure S5. Unsupervised clustering analysis of lung endothelial cells during LCMV infection.**

C57BL/6-F2r$^{fl/fl}$ mice were infected with LCMVArm strain and lung cells were isolated from naïve and infected mice at day 6 and 21 p.i and analyzed by flow cytometry expression of CD31, PDL1, tdTomato, CD34, CD309, VCAM-1, CD105, CD90.2 and CD201. 10,000 of manually gated lung endothelial cells (CD326$^-$CD45$^-$CD31$^+$) were concatenated from each naïve mice and infected mice at day 6 and 21 p.i. Unsupervised clusters were analyzed with Phenograph (k= 100). Heatmap of mean fluorescence intensity of each marker for all the clusters and bubble plots of median frequency of each cluster from naïve and infected mice (Figure 3). Frequency of each cluster is represented by box and whisker showing the median value with first and third quartiles in the box, with whiskers extending to the minimum and maximum values. Statistical significance was performed using non-parametric Mann-Whitney test. *P* value <0.05 was considered significant.

**Figure S6. Relationship between plasma levels of GZMA and T cell derived cytokines.**

**(A)** Relationship between plasma levels of CD8 T cell cytokines (IFNγ, TNFα, sFasL and IL-2) and granzymes (granzyme A and B) in PWH (n= 25). **(B)** *In vitro* activation and cytokine secretion by HUVEC. 5,000 HUVEC were stimulated with 20 ng/mL of rhIFNγ and 0.5 ng/mL of rhTNFα overnight and cytokines were measured in the supernatants. Detection of soluble adhesion molecules sVCAM-1 and sICAM-1 and cytokines IL-6, VEGF, Angiopoietin-2, FGFb, and IL-8. **(C)** *In vitro* HUVEC activation with granzymes. HUVEC were stimulated overnight with 100 nM of rhGZMA and rhGZMK in s the positive control, HUVEC were incubated with 100 μM of PAR1 agonist peptide (TFLLR-NH2) and control peptide (RLLFT-NH2). IL-6, IL-8 and Ang-2 were measured in the supernatants. Data is represented by box and whisker showing the median value with first and third quartiles in the box, with whiskers extending to the minimum and maximum values. Statistical significance was performed using non-parametric Mann-Whitney test. *P* value

<0.05 was considered significant. Correlations were performed using nonparametric Spearman correlation and $p$ value $\leqslant$ 0.01 was considered significant.

**Figure S7. PAR1 dependent activation by GZMA and GZMK.** HUVEC were stimulated overnight with 100 nM of rhGZMA and rhGZMK in the presence or absence of 10 μg/mL of ATAP-2 mAb or 300 nM of PAR1 antagonist SCH530348. As the positive control, HUVEC were incubated with 100 μM of PAR1 agonist peptide (TFLLR-NH2) and control peptide (RLLFT-NH2). VEGF, bFGF, sVCAM-1, sICAM-1 and sPCAM were measured in the supernatants. Data is represented by box and whisker showing the median value with first and third quartiles in the box, with whiskers extending to the minimum and maximum values. Statistical significance was performed using non-parametric Mann-Whitney test. $P$ value <0.05 was considered significant.

#### Table S1. Mouse Flow cytometry panels

#### Flow cytometry panel-1 (Endothelial cells staining)

| Specificity | Fluorochrome | Clone | Cat. # | Manufacturer |
|---|---|---|---|---|
| CD90.2 | BV570 | 30-H12 | 105329 | BioLegend |
| CD34 | PerCP-Cy5.5 | MEC14.7 | 119327 | BioLegend |
| CD105 | Pacific Blue | MJ7/18 | 120412 | BioLegend |
| CD326 | BV510 | G8.8 | 118231 | BioLegend |
| CD31 | BV605 | 390 | 102427 | BioLegend |
| CD45 | BV711 | 53-6.7 | 103147 | BioLegend |
| PD-L1 | BV785 | 10F.9G2 | 124331 | BioLegend |
| VCAM-1 | FITC | 429 (MVCAM.A) | 105706 | BioLegend |
| CD309 | PE-Vio770 | REA1116 | 130-119-436 | Miltenyi Biotech |
| CD16/CD32 | N/A | 2.4G2 | 553141 | BD Bioscience |

#### Flow cytometry panel-2 (Phenotypic characterization of lymphoid organs)

| Specificity | Fluorochrome | Clone | Cat. # | Manufacturer |
|---|---|---|---|---|
| B220 | BV510 | RA3-6B2 | 103247 | Biolegend |
| CD4 | BV605 | RM4-5 | 100547 | Biolegend |
| MHC II | BV655 | M5/114.15.2 | 107641 | Biolegend |
| CD45 | BV711 | 30-F11 | 103147 | Biolegend |
| CD11c | BV750 | N418 | 117357 | Biolegend |
| Gr-1 | FITC | RB6-8C5 | 108406 | Biolegend |
| CD3 | BUV805 | 17A2 | 569192 | BD |
| CD11b | BUV737 | M1/70 | 564443 | BD |
| CD8a | BUV496 | 53-6.7 | 569181 | BD |
| NK 1.1 | AF700 | S17016D | 156512 | Biolegend |
| CD16/CD32 | N/A | 2.4G2 | 553141 | BD Bioscience |

#### Flow cytometry panel-3 (LCMV-specific T cell staining)

| Specificity | Fluorochrome | Clone | Cat. # | Manufacturer |
|---|---|---|---|---|
| CD107a | BV785 | 1D4B | 121641 | BioLegend |
| CD45 | V570 | 30-F11 | 103136 | BioLegend |
| CD31 | V605 | 390 | 102427 | BioLegend |
| CD127 | BV650 | A7R34 | 135043 | BioLegend |
| CD44 | BV750 | IM7 | 103079 | BioLegend |
| CD3 | BUV805 | 17A2 | 569192 | BD Bioscience |
| CD8a | BUV496 | 53-6.7 | 569181 | BD Bioscience |
| KLRG1 | PE-Cy7 | 2F1/KLRG1 | 138416 | BioLegend |
| CD16/CD32 | N/A | 2.4G2 | 553141 | BD Bioscience |
| IFN$\gamma$ | APC | XMG1.2 | 562018 | BD Bioscience |
| TNF$\alpha$ | PerCP/Cy5.5 | MP6-XT22 | 506322 | Biolegend |
| GZMA | eFluor450 | 3G8.5 | 48-5831-82 | eBioscience |

#### Table S2. Flow cytometry panels

#### Flow cytometry panel-1 (PBMC staining)

| Specificity | Fluorochrome | Clone | Cat. # | Manufacturer |
|---|---|---|---|---|
| CD31 | BUV395 | 133.1 | 745600 | BD Bioseceince |
| CD4 | BUV805 | SK3 | 564910 | BD Bioseceince |
| CD3 | BV510 | UCHT1 | 563109 | BD Bioseceince |
| CD8 | BV605 | SK1 | 564115 | BD Bioseceince |
| CD49f | BV650 | GoH3 | 563706 | BD Bioseceince |
| PAR1 | PE | WEDE15 | IM2584 | Beckmen Coulter |
| CXCR4 | APC | 12G5 | 560936 | BD Bioseceince |
| CX3CR1 | APC-CY7 | 2A9-1 | 341615 | Biolegend |
| GZMA | AF700 | CB9 | 507210 | Biolegend |
| GZMB | BV421 | GB11 | 563389 | BD Bioseceince |

#### Flow cytometry panel-2 (sorting HIV-specific CD8 T cells)

| Specificity | Fluorochrome | Clone | Cat. # | Manufacturer |
|---|---|---|---|---|
| CD31 | PE-Dazzle 594 | WM59 | 303130 | Biolegend |
| CD45RA | V605 | HI100 | 562886 | BD Bioseceince |
| CD27 | B515 | M-T271 | 564642 | BD Bioseceince |
| CD3 | V650 | UCHT1 | 300468 | Biolegend |
| CD8 | PE-Cy7 | SK1 | 344750 | Biolegend |
| CXCR4 | APC | 12G5 | 560936 | BD Bioseceince |
| CD137 | PE | 4B4-1 | 561701 | BD Bioseceince |

Figure S1

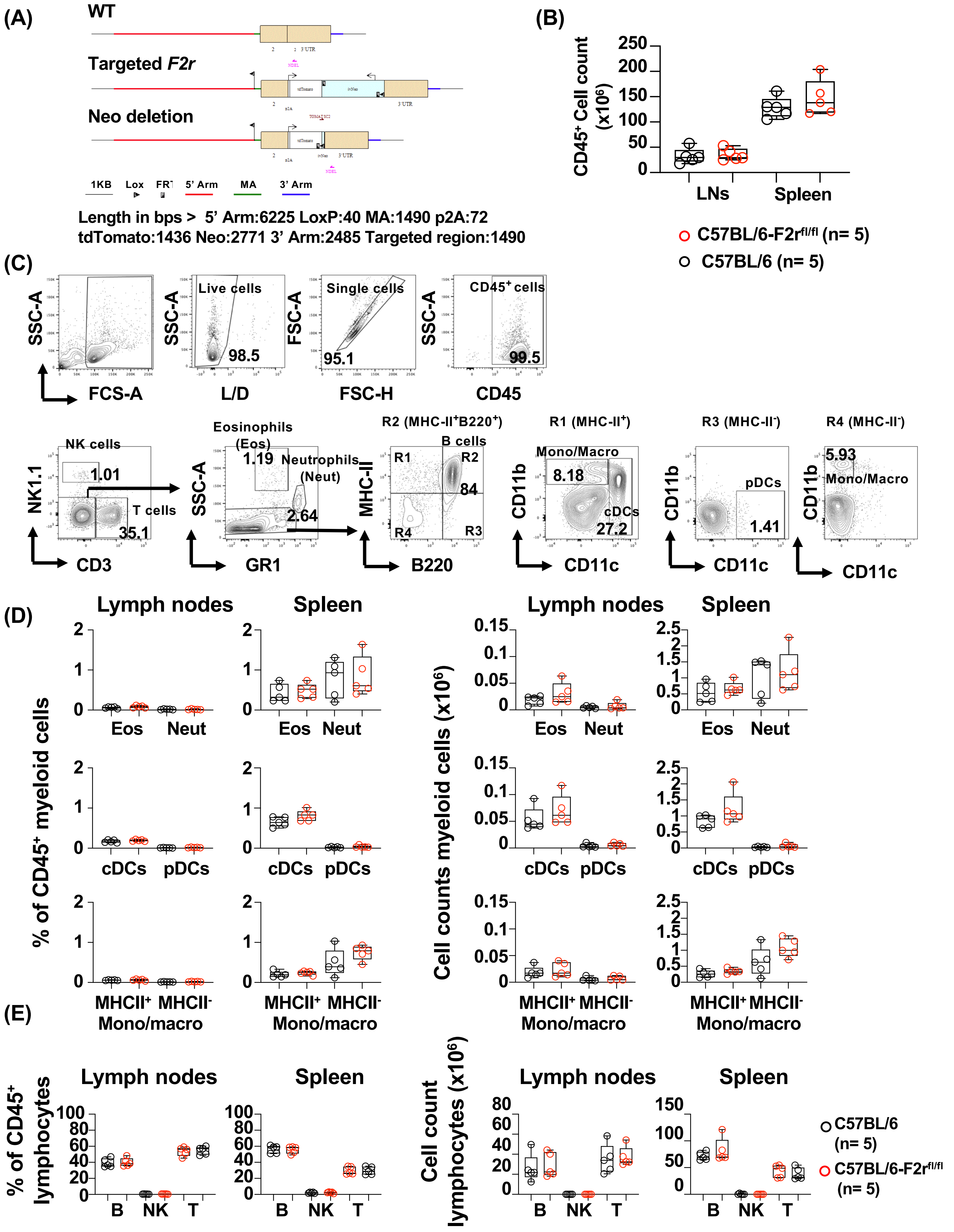
(A)
WT
Targeted F2r
Neo deletion
1KB Lox FRT 5' Arm MA 3' Arm
Length in bps > 5' Arm:6225 LoxP:40 MA:1490 p2A:72
tdTomato:1436 Neo:2771 3' Arm:2485 Targeted region:1490
(B)
CD45+ Cell count (x10^6)
LNs Spleen
C57BL/6-F2rfl/fl (n= 5)
C57BL/6 (n= 5)
(C)
Live cells 98.5
Single cells 95.1
CD45+ cells 99.5
FCS-A L/D FSC-H CD45
NK cells 1.01
T cells 35.1
Eosinophils (Eos) 1.19
Neutrophils (Neut) 2.64
R2 (MHC-II+B220+)
B cells 84
R1 (MHC-II+)
Mono/Macro 8.18
cDCs 27.2
R3 (MHC-II-)
pDCs 1.41
R4 (MHC-II-)
Mono/Macro 5.93
CD3 GR1 B220 CD11c CD11c CD11c
(D)
Lymph nodes Spleen
% of CD45+ myeloid cells
Cell counts myeloid cells (x10^6)
Eos Neut
cDCs pDCs
MHCII+ MHCII-
Mono/macro
(E)
% of CD45+ lymphocytes
Cell count lymphocytes (x10^6)
B NK T
C57BL/6 (n= 5)
C57BL/6-F2rfl/fl (n= 5)

### Figure S2

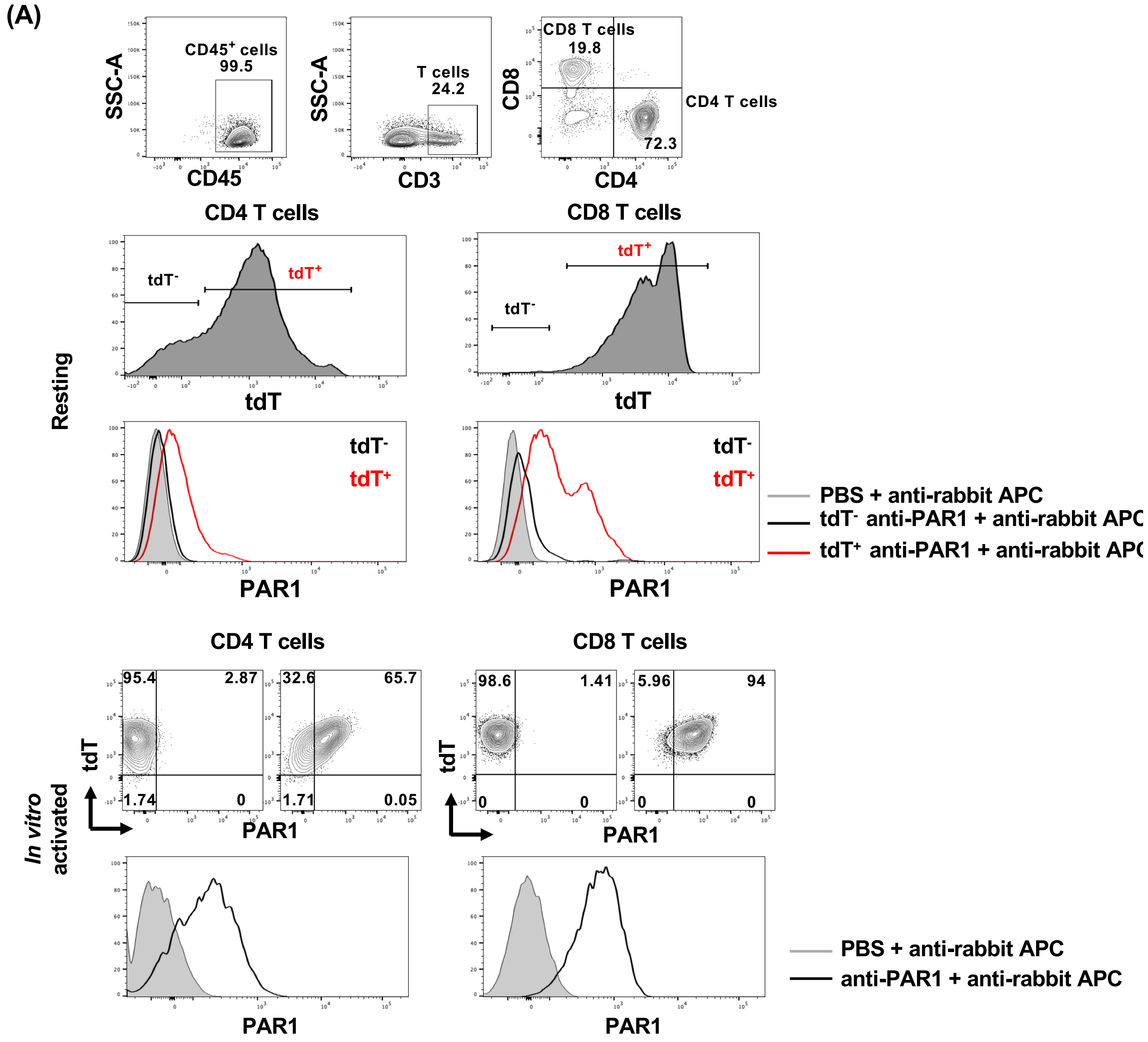
(A)
SSC-A
CD45+ cells
99.5
CD45
SSC-A
T cells
24.2
CD3
CD8
CD8 T cells
19.8
CD4 T cells
72.3
CD4
CD4 T cells
CD8 T cells
Resting
tdT-
tdT+
tdT
tdT-
tdT+
tdT
tdT-
tdT+
PAR1
tdT-
tdT+
PAR1
PBS + anti-rabbit APC
tdT- anti-PAR1 + anti-rabbit APC
tdT+ anti-PAR1 + anti-rabbit APC
CD4 T cells
CD8 T cells
In vitro activated
tdT
95.4
2.87
1.74
0
32.6
65.7
1.71
0.05
PAR1
tdT
98.6
1.41
0
0
5.96
94
0
0
PAR1
PAR1
PAR1
PBS + anti-rabbit APC
anti-PAR1 + anti-rabbit APC

Figure S3

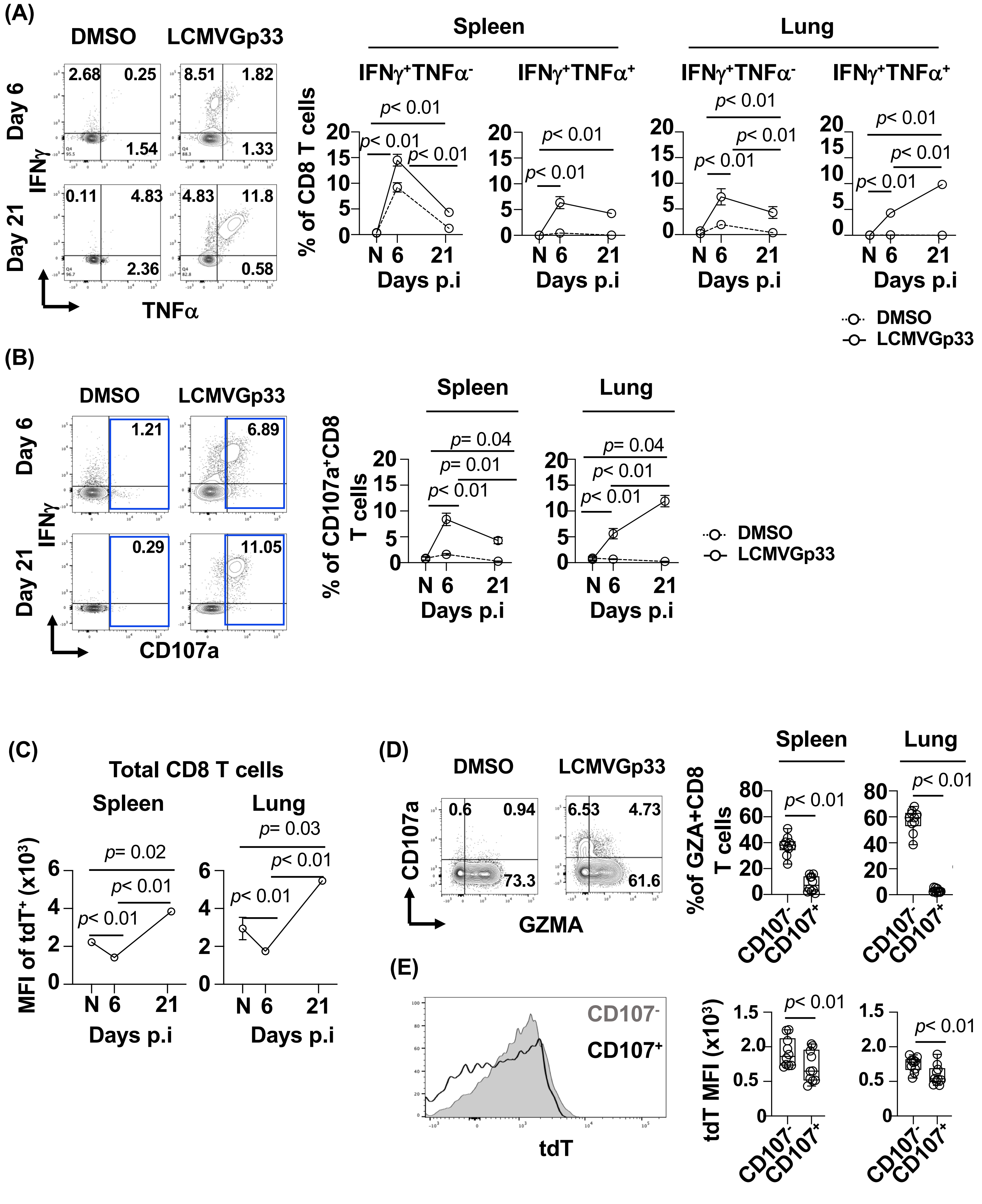

(A)
DMSO
LCMVGp33
Day 6
Day 21
IFNγ
TNFα
2.68
0.25
1.54
8.51
1.82
1.33
0.11
4.83
2.36
4.83
11.8
0.58
Spleen
Lung
IFNγ+TNFα-
IFNγ+TNFα+
% of CD8 T cells
Days p.i
DMSO
LCMVGp33
(B)
CD107a
1.21
6.89
0.29
11.05
% of CD107a+CD8 T cells
p= 0.04
p= 0.01
p< 0.01
(C)
Total CD8 T cells
MFI of tdT+ (x10³)
p= 0.02
p= 0.03
(D)
GZMA
0.6
0.94
73.3
6.53
4.73
61.6
%of GZA+CD8 T cells
CD107-
CD107+
(E)
tdT
tdT MFI (x10³)

Figure S4

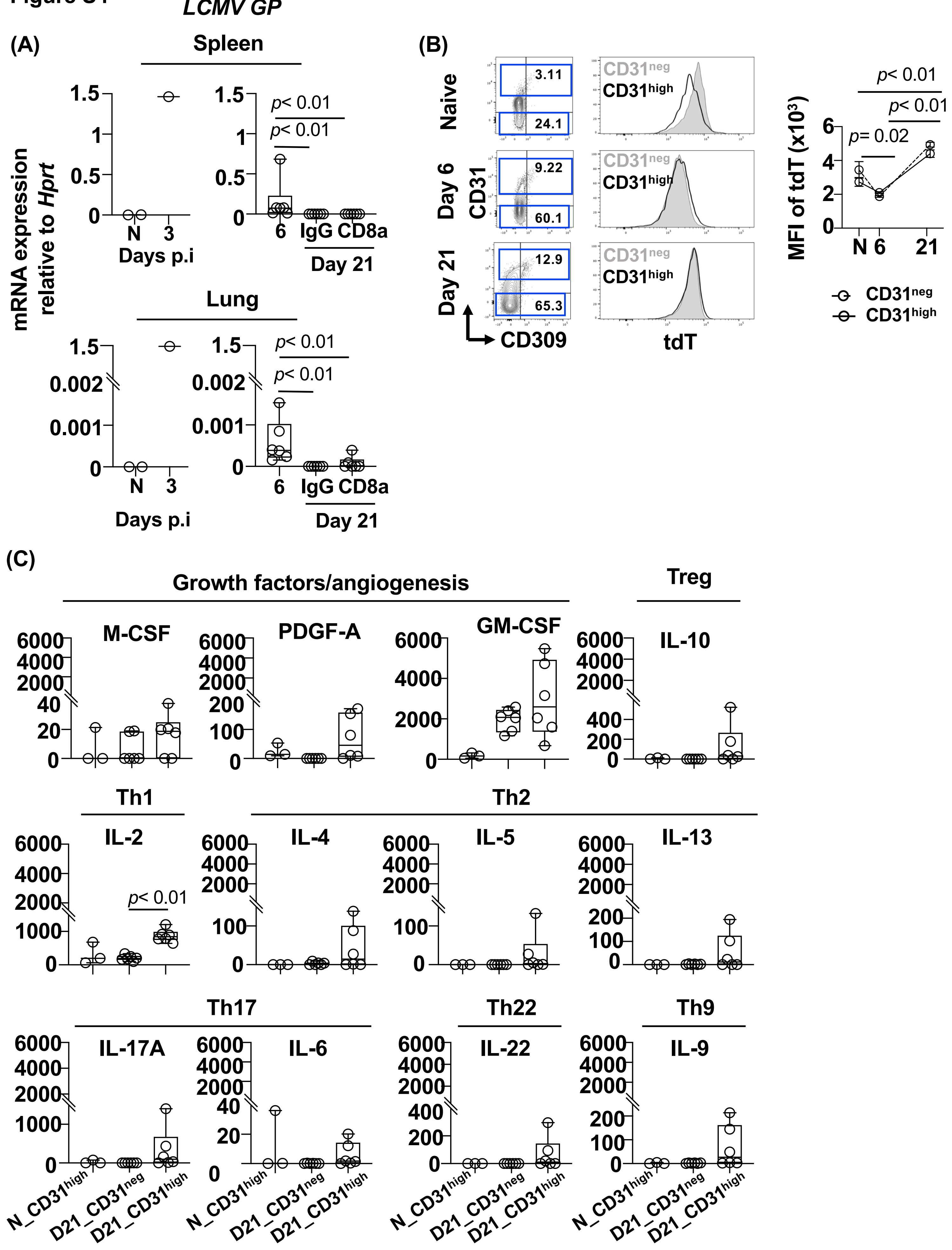
LCMV GP
(A)
Spleen
mRNA expression relative to Hprt
Days p.i
Day 21
p< 0.01
Lung
(B)
Naive
Day 6
Day 21
CD31
CD309
tdT
CD31neg
CD31high
MFI of tdT (x10³)
p= 0.02
(C)
Growth factors/angiogenesis
M-CSF
PDGF-A
GM-CSF
Treg
IL-10
Th1
IL-2
Th2
IL-4
IL-5
IL-13
Th17
IL-17A
IL-6
Th22
IL-22
Th9
IL-9
N_CD31high
D21_CD31neg
D21_CD31high

#### Figure S5

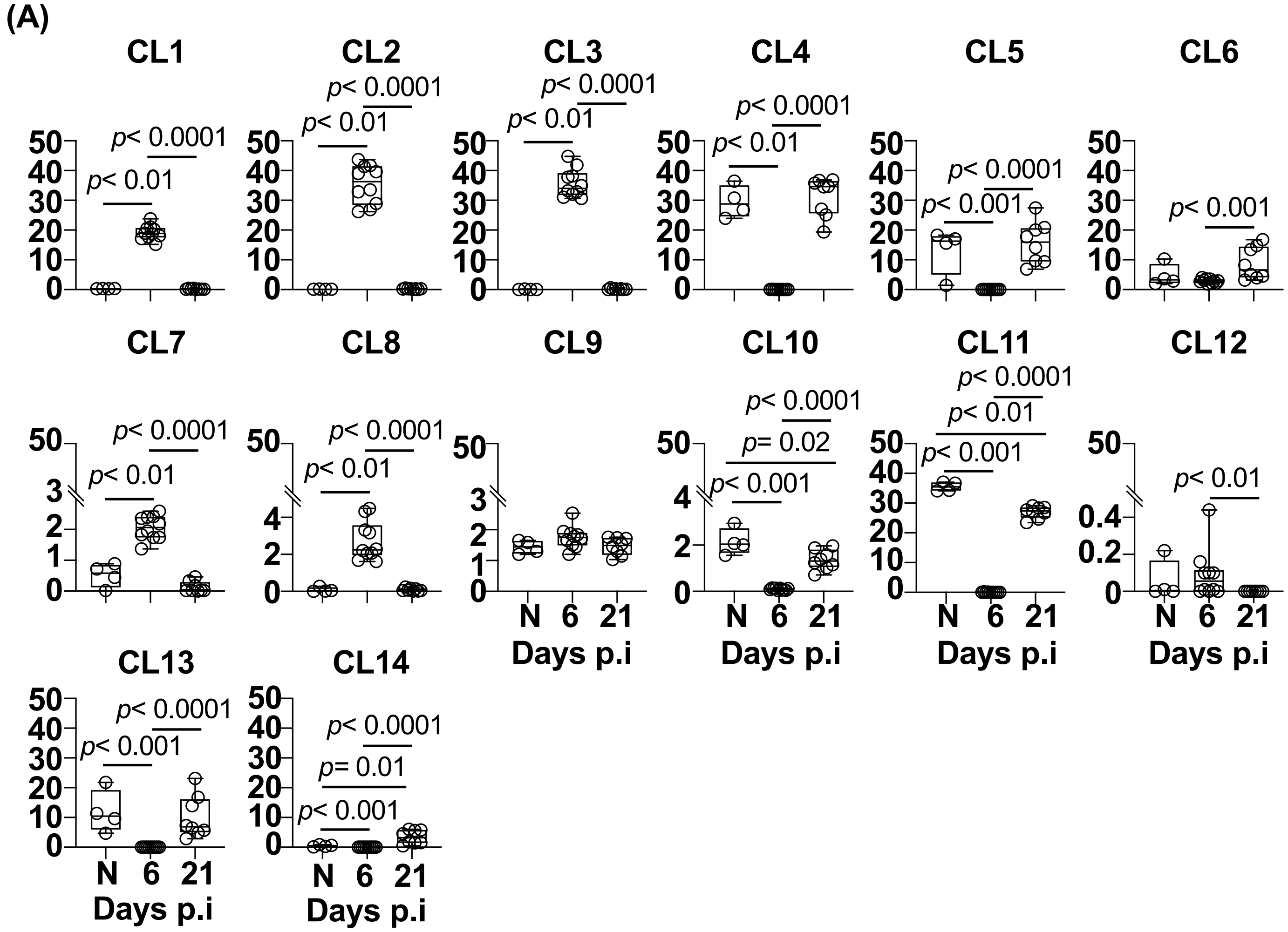
(A)
CL1
CL2
CL3
CL4
CL5
CL6
CL7
CL8
CL9
CL10
CL11
CL12
CL13
CL14
p< 0.0001
p< 0.01
p< 0.001
p= 0.02
p= 0.01
N
6
21
Days p.i

Figure S6

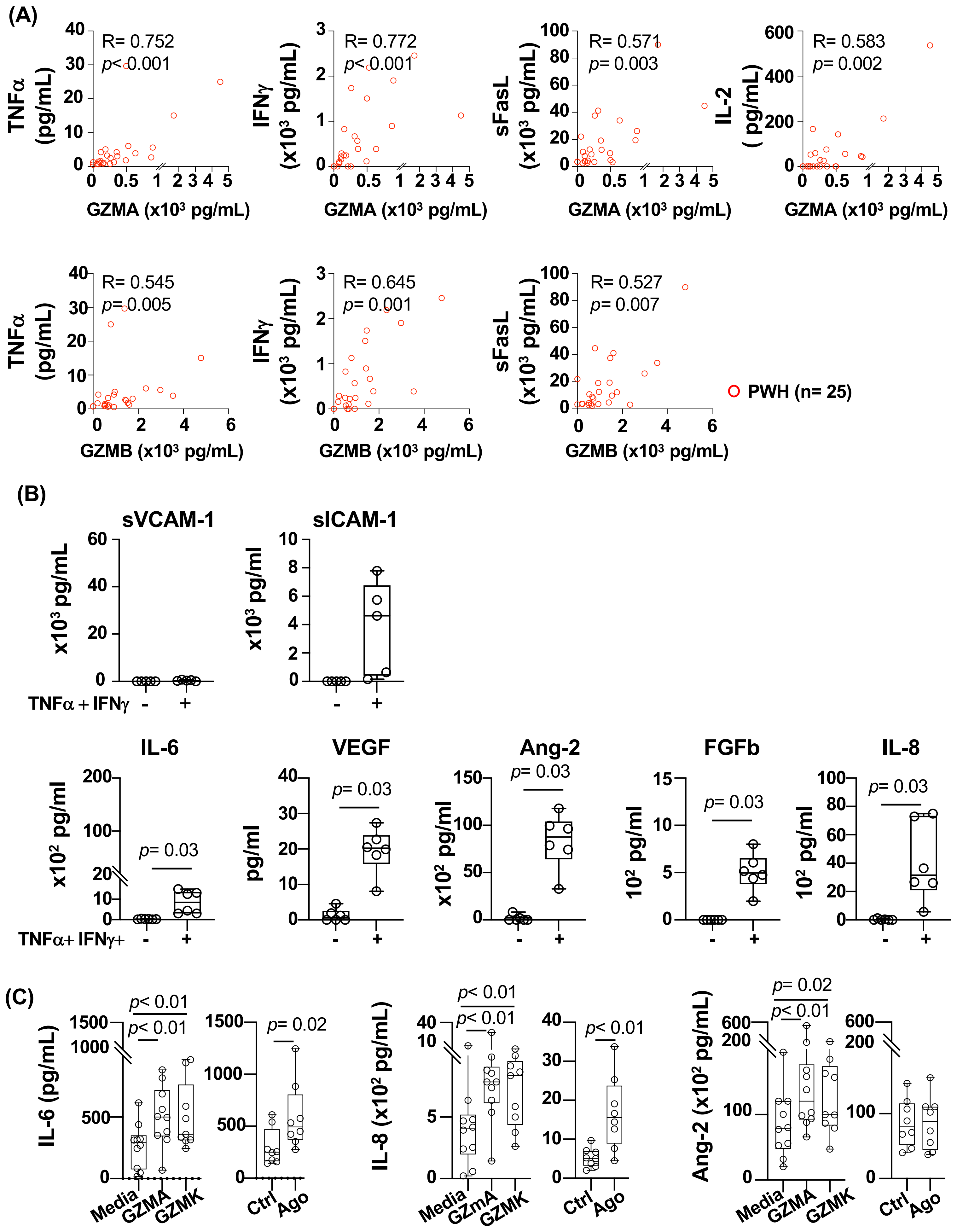

(A)
R= 0.752
p< 0.001
TNFα (pg/mL)
GZMA (x10³ pg/mL)
R= 0.772
p< 0.001
IFNγ (x10³ pg/mL)
GZMA (x10³ pg/mL)
R= 0.571
p= 0.003
sFasL (x10³ pg/mL)
GZMA (x10³ pg/mL)
R= 0.583
p= 0.002
IL-2 ( pg/mL)
GZMA (x10³ pg/mL)
R= 0.545
p= 0.005
TNFα (pg/mL)
GZMB (x10³ pg/mL)
R= 0.645
p= 0.001
IFNγ (x10³ pg/mL)
GZMB (x10³ pg/mL)
R= 0.527
p= 0.007
sFasL (x10³ pg/mL)
GZMB (x10³ pg/mL)
PWH (n= 25)
(B)
sVCAM-1
x10³ pg/mL
sICAM-1
x10³ pg/ml
TNFα + IFNγ - +
IL-6
x10² pg/ml
p= 0.03
VEGF
pg/ml
p= 0.03
Ang-2
x10² pg/ml
p= 0.03
FGFb
10² pg/ml
p= 0.03
IL-8
10² pg/ml
p= 0.03
TNFα+ IFNγ+ - +
(C)
IL-6 (pg/mL)
p< 0.01
p< 0.01
Media GZMA GZMK
p= 0.02
Ctrl Ago
IL-8 (x10² pg/mL)
p< 0.01
p< 0.01
Media GZmA GZMK
p< 0.01
Ctrl Ago
Ang-2 (x10² pg/mL)
p= 0.02
p< 0.01
Media GZMA GZMK
Ctrl Ago

**Figure S7**

**(A)**

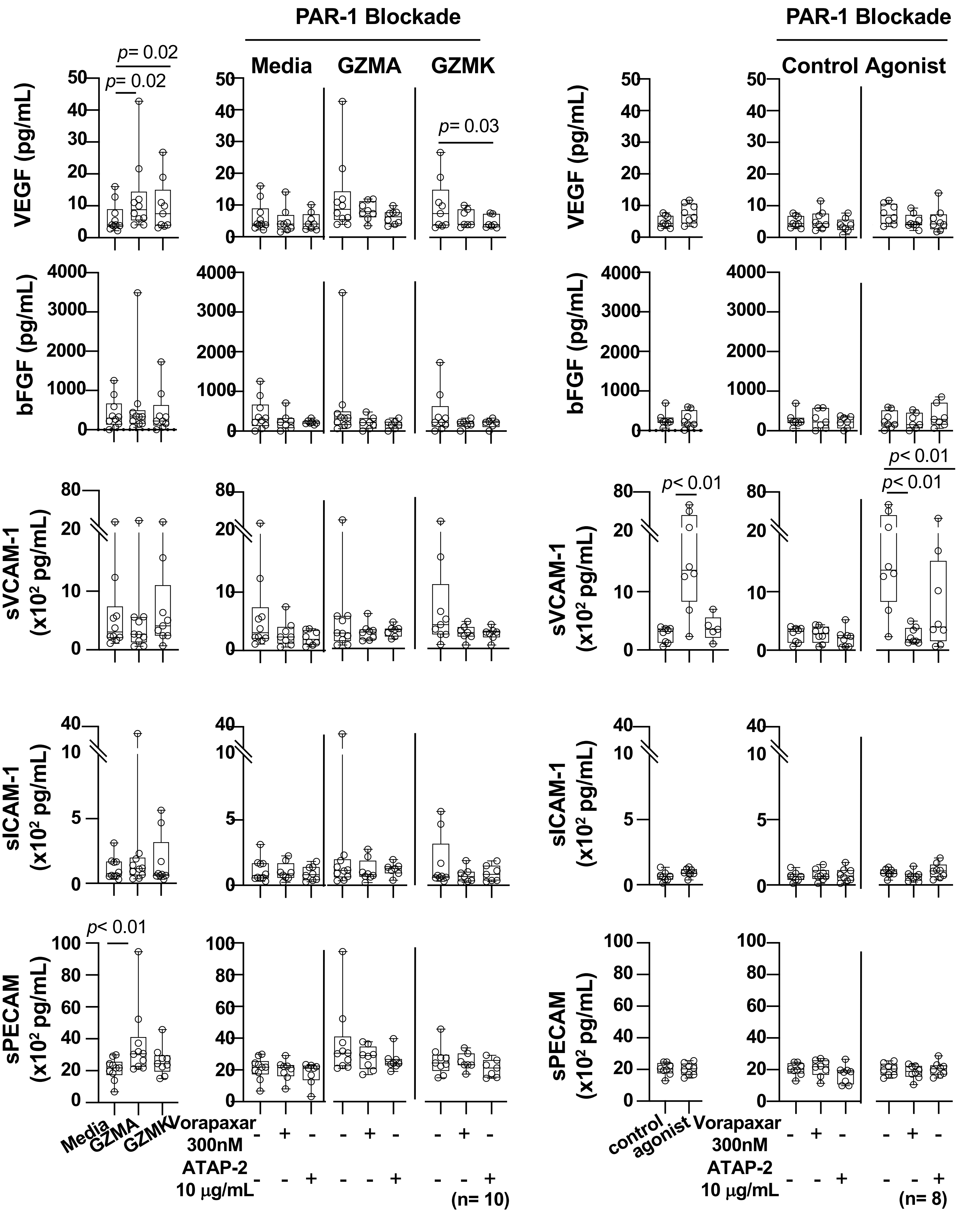
PAR-1 Blockade
PAR-1 Blockade
Media
GZMA
GZMK
Control Agonist
VEGF (pg/mL)
bFGF (pg/mL)
sVCAM-1 (x10² pg/mL)
sICAM-1 (x10² pg/mL)
sPECAM (x10² pg/mL)
p= 0.02
p= 0.02
p= 0.03
p< 0.01
p< 0.01
p< 0.01
p< 0.01
Media
GZMA
GZMK
control
agonist
Vorapaxar 300nM
ATAP-2 10 μg/mL
(n= 10)
(n= 8)
